# Enhanced local cooling effects of forests across the globe

**DOI:** 10.1101/2023.10.17.562656

**Authors:** William W. M. Verbiest, Gabriel Reuben Smith, Leila Mirzagholi, Thomas Lauber, Constantin M. Zohner, Daniel S. Maynard, Sebastian Schemm, Thomas W. Crowther

## Abstract

Forests cool the land surface in warm regions and warm the land surface in cool regions. Because these local temperature buffering effects depend on background climate^1,2^, increasingly large areas might experience forest cooling effects as the climate warms^3–5^. Here, using statistical modeling, we quantified changes in the sensitivity of land surface temperatures^6^ to forest cover change^7^ from 1988 – 2016, across 15 million km^2^ of land area. Forests had a local cooling effect on day and night surface temperatures in 86.3% (72 million km^2^) and 61.6% (5,108 million km^2^) of the forest area, respectively. This area has increased by 0.3% (0.2 million km^2^) at day and 4.7% (2 million km^2^) at night over recent decades. Our study indicates that climate change is enhancing the cooling effect of forests in the short term, underscoring the importance of protecting natural, diverse forest ecosystems in the face of rising global temperatures that threaten ecosystems^8,9^, human health^10,11^, and food production^12^.

## Main

Forests affect Earth’s climate system by serving as a major terrestrial carbon sink^13–15^ and affecting local temperatures through biophysical processes^16–18^. The conservation and restoration of natural forests are therefore key components in the global effort to fight climate change^19,20^. However, it remains unclear how the biophysical effects of forests are changing in a warming climate, which limits our capacity to predict the climatic impacts of forests across the globe under future climate change^21–23^.

The biophysical effects of forests on local temperatures are strongly determined by the background climate, including temperature, rainfall, and snow cover^3–5^. This creates latitudinal patterns in forests’ local climatic impacts whereby forests have a cooling effect in warm, tropical regions and a warming effect in colder, high-latitude regions^17,23–25^. Given this large influence of background climate on how forests affect local temperatures^3–5^, climate change is likely to fundamentally alter these patterns^5,26^, with the potential to enhance the cooling effects of forests as the planet warms^3,5^. Yet, the extent and direction of changes in these effects due to climate change remain poorly understood at a global scale^5^, limiting our capacity to assess the climate impacts of policy measures such as forest conservation or restoration.

To test how changes in background climate are shifting the local temperature effects of forests across different latitudes and climates, we quantified changes in the sensitivity of land surface temperatures to forest cover change (ΔT/ΔFC) over the past three decades (1988-2016) using multivariate linear regression. To do so, we analysed MODIS land surface temperature data obtained from 650,928 pixels across 15 million km^2^ forest area that underwent significant forest cover changes (ΔFC > 2%) between 2003 and 2012 (refs^6,7^), whilst accounting for the influence of background climate^27,28^, topography^29^, and soil type^30^. By comparing these effects under past and present climate conditions^27,28,31^, we then estimated the change in the proportion of land where forests cool the Earth’s surface. Finally, to generate a more mechanistic understanding of how climate change might affect the sensitivity of land surface temperatures to forest cover, we quantified the drivers of the changes in the Earth’s surface energy balance by isolating the impact of changes in individual climate variables.

### Local surface temperatures and forest cover change

We found that a loss of forest cover resulted in increased mean surface temperatures (i.e., average of day and night temperature) in all biomes, except for boreal forest, where it led to surface cooling (Fig. 1a). This suggests that forests act as natural temperature regulators by buffering local temperatures, with a cooling effect in warmer regions and a warming effect in colder regions^1,2,8,32–35^. Arid regions showed the strongest changes in mean surface temperature in response to forest cover loss, followed by the tropical, temperate, and boreal biomes, with the latter region getting cooler as forest cover decreases (Table S2). This pattern can be explained by the effect of incident solar radiation, which decreases with latitude and cloudiness^32^. Indeed, the response of mean surface temperatures to forest cover changes followed a clear latitudinal gradient, with a response shift from positive to negative occurring at around 50°N (Fig. 1b).

**Fig. 1:**
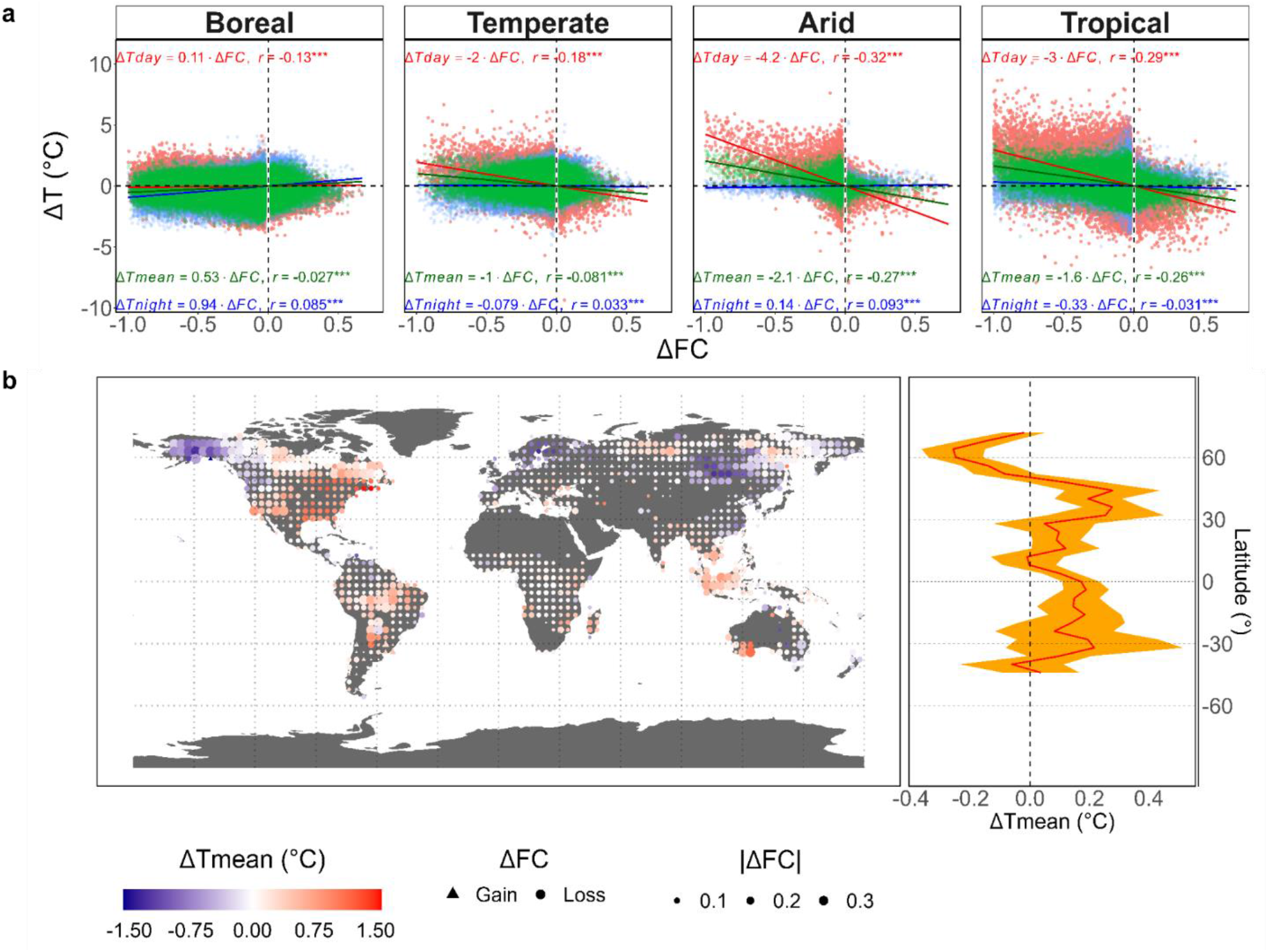
Surface temperature response to forest cover change for 2003-2012. **a**, Changes in day-time, mean (i.e., average of day and night response), and night-time surface temperature (Δ*T*) as a function of forest cover change (Δ*FC*) for the boreal (n=195,892), temperate (n=158,851), arid, (n=17,598) and tropical biome (n=278,587). The coefficient of the univariate linear regression model (Δ*T* = 0 + *β* ∗ Δ*FC*) is shown alongside the Pearson correlation coefficient (r) and P-value. P-values correspond to <0.001 (***), <0.01 (**), <0.05 (*), <0.1 (.), and >0.1 (). **b,** Change in mean surface temperature (Δ*Tmean*) due to forest cover changes at 4° resolution (left) (n=1,014). The direction and magnitude of the forest cover changes are indicated by the shape and size of the symbols, respectively. The latitudinal pattern of Δ*Tmean* at 4° latitudinal interval (right) (n=1,009). Latitudinal intervals with less than 3 data points were excluded. The 95% confidence interval is shown in orange.

We also evaluated the effects of forest cover on day and night surface temperatures and found similar changes (Fig. 1a). Day surface temperatures in arid, tropical, and temperate biomes tended to increase as forest cover decreased, while the response of day surface temperature to forest cover change in boreal regions was negligible (Table S2). Conversely, night surface temperatures showed a different pattern, with forest loss leading to cooling in boreal regions and warming in the other biomes (Table S2).

### The background climate and local surface temperature effects of forests

Through analysing the contribution of the Earth’s surface energy balance to the effect of forest cover change on local surface temperatures, we show that the biophysical effects of forest cover change are determined by background climate conditions in each biome and climate zone, thus confirming previous studies^17,23–25^ (Fig. S5; Fig. S6). In warmer, low-latitude regions, the non-radiative effects of forest cover change (i.e., heat fluxes) are more significant, while in colder, high-latitude regions, changes in radiative fluxes (i.e., reflected shortwave radiation) become more important due to the high reflectance of snow (Fig. S1b; Fig. S5; Fig. S6)^5,36^.

To demonstrate the effect of the background climate on local temperature changes due to forest cover change, we then isolated the individual effects of the climate variables in the statistical models, i.e. mean annual temperature, mean precipitation of driest quarter, and sum of snow water equivalent, on temperature responses to forest cover change. Our results show that mean annual temperatures substantially modulate temperature responses to forest cover change (Fig. 2), with forest loss resulting in surface warming during the day, particularly in warm regions with high mean annual temperatures. Similarly, forest loss resulted in surface warming during the night in warm regions but had a cooling effect in colder locations (Fig. 2). In snow-covered regions, forest loss led to surface cooling due to higher albedo reflectance (Fig. S7).

**Fig. 2:**
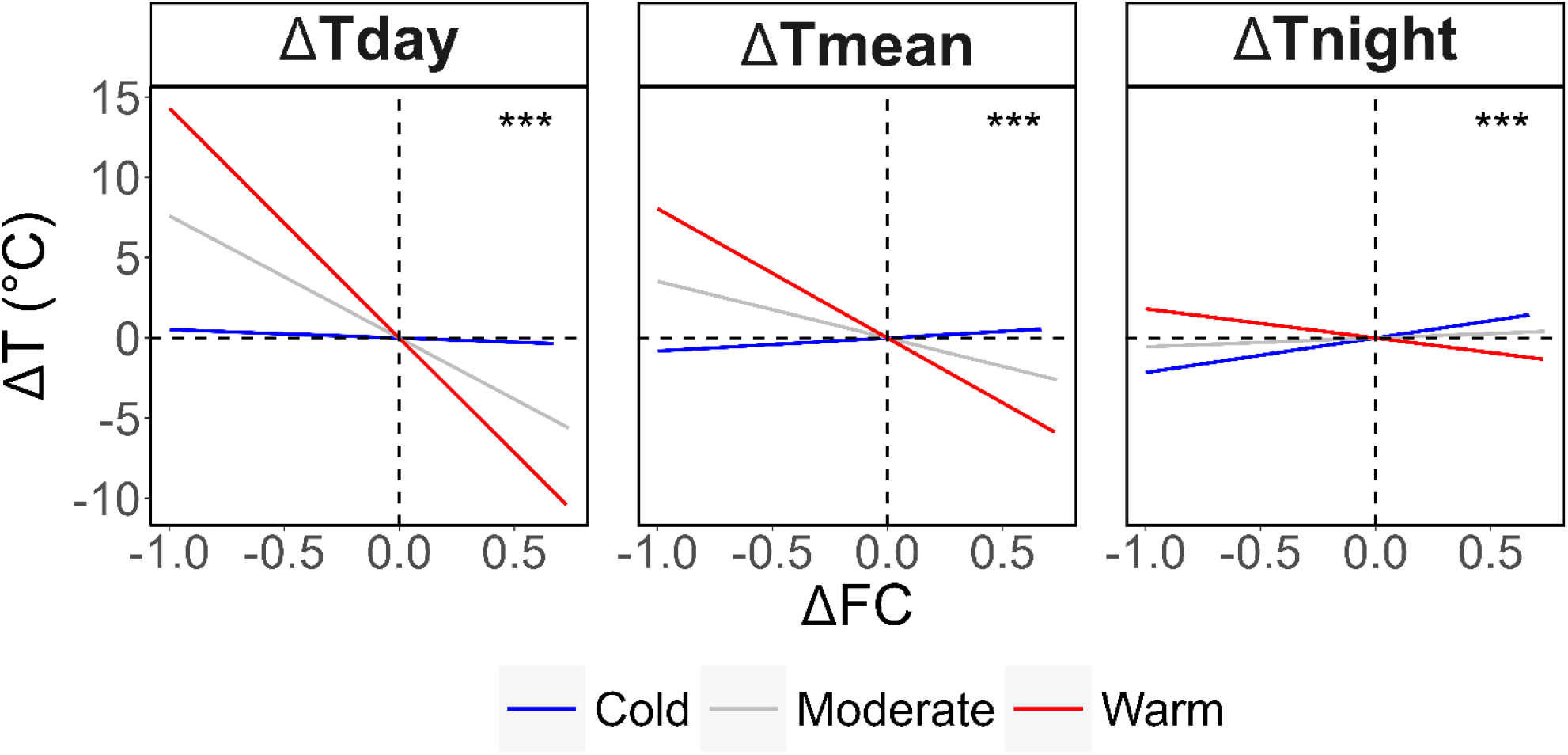
The interaction between mean annual temperatures and the sensitivity of surface temperatures to forest cover change. Biome-level model partial residuals in respect of the interaction between forest cover change and mean annual temperatures for the models predicting day, mean (i.e., average of day and night response), and night surface temperature (Δ*T*) change as a function of forest cover change (Δ*FC*). The cold (blue; n=65,093), moderate (grey; n=520,756), and warm (red; n=65,079), which corresponds to 0-0.1, 0.1-0.9, and 0.9-1 quantile intervals of the average mean annual temperature of the period 2003-2012, respectively. Global-level linear regression was performed for each climate zone. The stars indicate the significance of the interaction between mean annual temperature and Δ*FC* from a global level combining all climate zones.

### Changes in local cooling effects of forests over the past three decades

To quantify how climate change has affected the impact of forest cover changes on land surface temperatures over the past three decades, we combined our biome-level models with annual climate data from 1988 to 2016^27,28,31^. The results showed that globally, the ability of forests to cool the surface has increased over the past three decades by 3.9% (0.03°C per 100% forest restoration) for mean surface temperatures (Fig. S11). This is because the decrease in surface cooling due to latent heat fluxes was outweighed by an increase in surface cooling due to sensible heat fluxes and a decrease in surface warming due to shortwave reflected radiation (Fig. S11). In most low-latitude regions, the cooling effect of forests increased due to enhanced sensible heat fluxes (Fig. 3a; Fig. S14). However, in many areas, this cooling effect was partly offset by a reduction in latent heat fluxes under increased mean annual temperatures (Fig. S14). This might correspond with a potential increase in the water-use efficiency (i.e., similar photosynthesis rate at a lower plant transpiration rate) over the past few decades due to CO_2_ fertilization^37,38^.

**Fig. 3:**
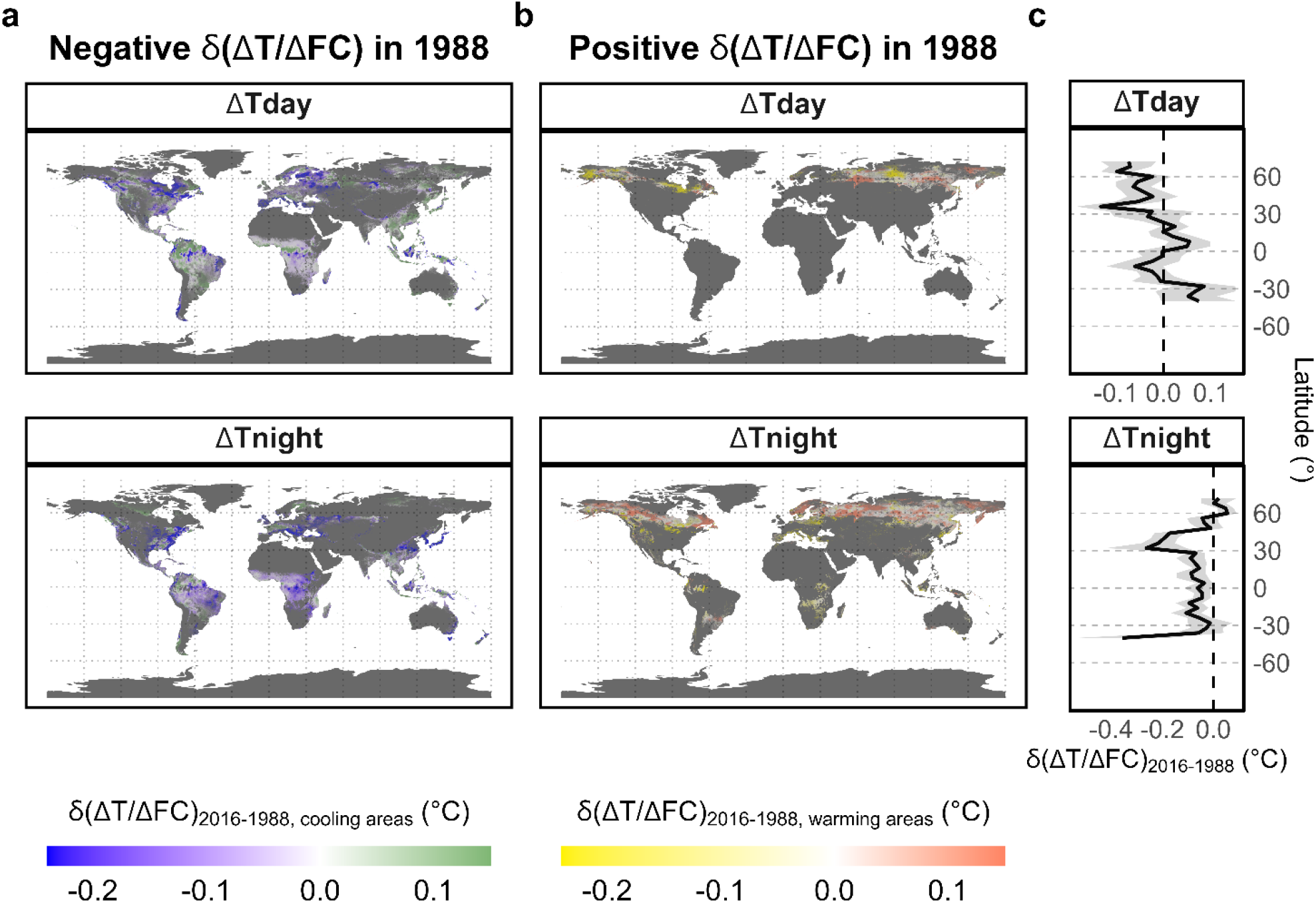
Change in sensitivity of surface temperatures to forest cover change for 1988-2016. **a,** Changes in the sensitivity of the day and night surface temperature to forest cover change (δ(Δ*T*/Δ*FC*)) between 1988 and 2016 for pixels with a negative Δ*T*/Δ*FC* in 1988 (i.e., surface cooling with forest gain; n=2,719,786). Blue (green) indicates an increase (decrease) in the cooling effect. **b,** Change in the sensitivity of the day and night surface temperature to forest cover change (δ(Δ*T*/Δ*FC*)) between 1988 and 2016 for pixels with a positive Δ*T*/Δ*FC* in 1988 (i.e., surface warming with forest gain; n=1,044,226). Yellow (red) indicates a decrease (increase) in the warming effect. **c,** Latitudinal pattern of the difference in the sensitivity of day and night surface temperatures to forest cover change (δ(Δ*T*/Δ*FC*)) between 1988 and 2016 based on 4° latitudinal means. Latitudinal intervals with less than 3 data points were excluded. The 95% confidence interval is shown in grey.

At higher latitudes, temporal changes in the warming effects of forests showed high spatial heterogeneity (Fig. 3b-c). Our statistical models show that regions where snowfall has decreased over the past three decades have experienced a decrease in the warming effect of forests due to a reduction in shortwave reflected radiation in non-forest areas (Fig. 3b; Fig. S10b; Fig. S13; Fig. S14). This decrease in snow cover has narrowed the temperature difference between forested and non-forested land, resulting in a decrease in the warming effect of forests. Furthermore, an increase in mean annual temperatures has led to extensions in the growing season at higher latitudes^39^, which is likely to result in increased evapotranspiration rates and latent heat fluxes (Fig. S11; Fig. S13; Fig. S14). This process helps to cool the surrounding area, offsetting the warming effect of forests in those regions and potentially even reversing it to a cooling effect in some places.

To determine the spatial distribution of these temporal changes in forests’ cooling effects, we calculated the annual proportion of the area where forests caused surface cooling compared to the total forested area from 1988 to 2016. In 1988, 86.3% (72 million km^2^) and 61.6% (51 million km^2^) of forested regions showed cooling during the day and night, respectively, mainly at lower and mid-latitudes (Fig. 3a). Over the next three decades, the global proportion of land where forests had a cooling effect increased by 0.3% (0.2 million km^2^) during the day and 4.7% (2.4 million km^2^) during the night (Fig. 4).

**Fig. 4:**
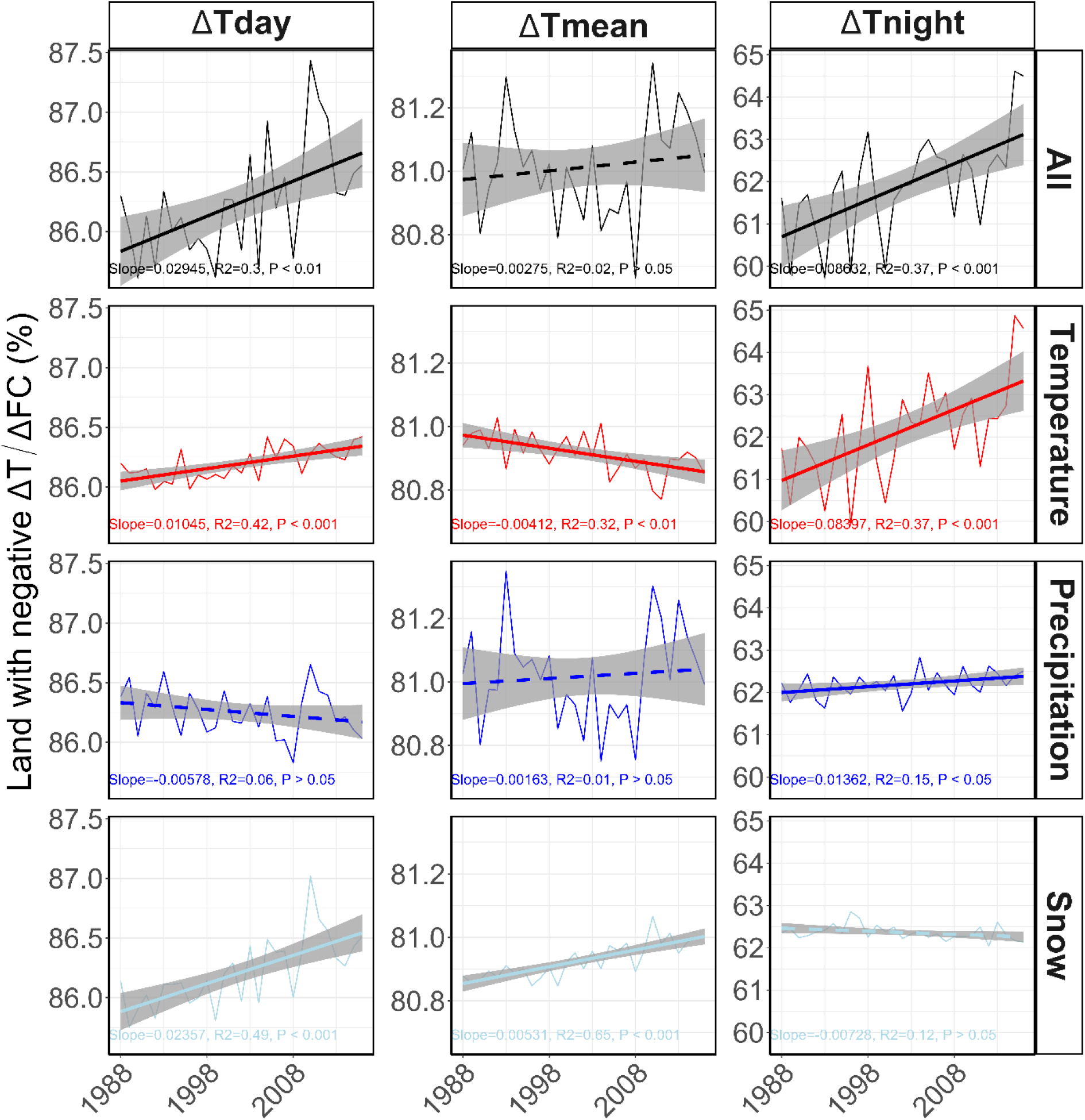
Temporal change and climatic drivers of the sensitivity of surface temperatures to forest cover change (Δ*T*/Δ*FC*) between 1988 and 2016. The temporal change of the proportion of land with a negative Δ*T*/Δ*FC* was calculated for the day, mean (i.e., average of day and night response), and night Δ*T*/Δ*FC* for the period 1988-2016 (n=3,764,012). Changes in precipitation (mean precipitation of driest quarter), snow (sum of snow water equivalent), and temperature (mean annual temperature) were separately assessed while other climate variables were kept constant and compared to the scenario in which all climate variables change over time (All). Solid lines represent regression lines with significant P-values (<0.05), while dashed lines denote non-significant regression lines (P-values >0.05).

The shift from a warming to a cooling effect of forests was more pronounced for night-time temperatures at lower and mid-latitudes, while for day-time temperatures, this shift was mainly observed in boreal regions (Fig. S15). As a result, the positive alteration in the proportion of land where forests cool the surface was almost three times greater for night-time temperatures (0.086 ± 0.022 %/year, R^2^ = 0.37, P-value < 0.001) compared to day-time temperatures (0.029 ± 0.009 %/year, R^2^ = 0.3, P-value < 0.01) (Fig. 4). This increase in the cooling effect of forests is due to both an increase in mean annual temperatures and a reduction in snow cover (Fig. 4; Fig. S11; Fig. S14) and is expected to continue as global change intensifies^3,5^.

Despite confidence in our estimates of local surface temperature sensitivity to forest cover changes and their relation to background climate (Fig. S2; Fig. S12), several assumptions in our approach contribute to uncertainty in our results. Firstly, we did not distinguish between the types of land cover transition, which can also affect local temperature changes, and contributed considerable noise to our dataset^2,40^. Secondly, our observations of forest cover change were based on a limited duration of ten years, which was the longest duration available at the time of our study.

Thirdly, the use of satellite imagery has drawbacks, including low sampling frequency and observational bias toward non-overcast conditions^36^. Fourthly, we acknowledge the uncertainties related to the satellite data products^41,42^. Finally, our observational approach cannot capture the impact of changes in forest cover on the hydrological cycle^12,33^, sea temperature-surface albedo interactions^17,43,44^, downwelling shortwave radiation^5^, and non-local climatic effects^44,45^, all of which are likely to contribute to the remaining unexplained variation in the observed forest temperature effects (Fig. 1; Fig. S16). Nevertheless, despite these uncertainties, our approach allowed us to reveal general underlying changes that reveal clear biophysical effects of forests at a high resolution. Our findings are based on statistical extrapolations from observations, and future research, using alternative approaches, such as climate models^2,21,46,47^ and *in situ* measurements^2,48–50^, can further increase our mechanistic understanding of these biophysical patterns.

Our study examines how the effect of forests on local surface temperature varies with background climate, highlighting the short-term consequences of climate change on how forests modulate the local climate. We show that forests generally have a local warming effect in high latitude regions, but a cooling effect in the majority of the global forest area (86.3% of the land surface in the day, and 61.6% at night). We also show that the local temperature effect of forests has shifted considerably over the past three decades, enhancing the cooling effects worldwide and increasing the global area where forests have cooling effects (by 0.2 million km^2^ in the day, and 2.4 million km^2^ at night). This short-term increase in the local cooling effect of forests was mainly driven by climate warming, which predicts that this cooling effect will become more pronounced as the planet continues to warm. Our results are integral to guiding future climate mitigation policies as they show that forest conservation will become increasingly important for cooling local surface temperatures across the globe. As we continue to lose forests worldwide, we remove their capacity to buffer the effects of climate warming^5,21^, creating negative feedbacks that are likely to drive further warming effects. These climate benefits of forests further underscore the urgency to halt and reverse the loss of diverse, natural forests worldwide, for the preservation of the well-being of local people and biodiversity^11,12^.

## Methods

### Data preparation and processing

#### Forest cover and land surface temperatures

Satellite-derived information of changes in forest cover (ΔFC)^7^ and local land surface temperature (ΔT)^6^ were processed in Google Earth Engine^51^. To estimate ΔT as a response to ΔFC ^1^, we opted not to use the space-for-time logic. This approach implies that local temperature responses to land cover transitions are estimated by comparing temperatures of adjacent contrasting vegetation types^32,52,53^ since this approach gives biased estimates as forests are not randomly distributed across the landscape^1^. Instead, for the final analysis, we calculated the temperature change between 2003 and 2012, since a more robust climate response was assumed for the longest considered period^1^. We also examined the nine different year combinations between 2003-2005 and 2010-2012, and every year between 2004 and 2011 with 2003 as a baseline to assess the inter-annual variability of the temperature response to forest cover change and the performance of the models used for predicting surface temperature sensitivities to forest cover change (see below).

To estimate ΔFC, we used high-resolution global maps of 21^st^-century forest cover changes at one-arcsec resolution from Hansen et al.^7^. This data includes the forest cover in 2000 (ranging between 0-100%), the forest gains between 2000-2012, and the year in which gross forest cover loss occurred between 2000-2021. We chose to limit our study to the period during which data on forest gains are available, i.e. 2000-2012.We defined forest as pixels with a tree cover >10%, and a tree height >4 m^18^. To match ΔFC to the temperature data (see below), the maps were transformed from 1 arcsec (∼0.00028°) to 0.05° and reprojected in WGS84 (EPSG:4326) (see below).

To calculate ΔFC, we took the difference in forest cover of the two considered years with ΔFC ranging from −1 (100% deforestation) to 1 (100% forest gain). Observed forest gains were assumed to reflect linear increases over time, as in Alkama and Cescatti^1^, since these data are only available as a single measurement across the period of 2000-2012 ^7^. Forest cover changes between 2003 and 2012 mainly occurred in the tropics (mean ΔFC ± standard deviation: −0.09 ± 0.12; n = 278,587), followed by boreal (−0.11 ± 0.20; n =195,892), temperate (−0.07 ± 0.11; n = 158,851), and arid regions (−0.07 ± 0.13; n = 17,598).

To calculate ΔT, we used radiometric land surface temperatures (LST) measured by the Moderate Resolution Imaging Spectroradiometer MYD11C3 (MODIS) satellite^6^. ^1^We chose to use MODIS AQUA instead of MODIS TERRA data, since the former better matches the timing of the minimum and maximum surface temperature data ^1^. First, we acquired the daily maximum and minimum LST at 0.05° resolution. Then, we calculated the difference between the two focal years. The change in mean LST was calculated as the mean of the change in the day and night LST^1^.

The total temperature response (ΔT) consists of the temperature change due to forest cover changes (ΔT_fcc_) and a residual temperature change caused by natural climate variability (ΔT_res_). Using the inverse distance weighting method, we estimated Δ*T*_*res*_ for pixels with a significant ΔFC (ΔFC < −0.02 or ΔFC > 0.02) based on the ΔT of surrounding stable forests (−0.02 < ΔFC < 0.02 where ΔT_fcc_ ≅ 0 and thus ΔT ≅ ΔT_res_) within a radius of 50 kilometers^1^:

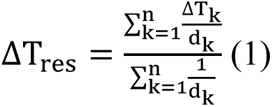

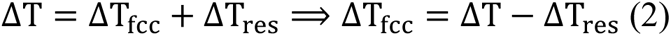

where *d* is the distance of the pixel with significant ΔFC to the pixel k containing stable forest. For the remaining analysis, we then used ΔT_fcc_and ΔFC at locations with a significant ΔFC.

#### The Earth’s surface energy balance

To understand which components of the Earth’s surface energy balance are responsible for the observed ΔT, we estimated changes in latent heat fluxes, the shortwave reflected radiation, the longwave emitted radiation, and the combined ground and sensible heat fluxes between 2003 and 2004-2012. We also did this for the nine different year combinations of 2003-2005 and 2010-2012.

First, we obtained yearly estimates for all data products by averaging the monthly values. The latent heat flux data (LE) came from the MODIS MOD16A2 product^54^ without modification. We calculated shortwave reflected radiation (SW_↑_) by multiplying the downwelling shortwave radiation (SW_↓_) from Kato et al.^55^ by the MODIS MCD43C3 surface albedo^6^. The surface albedo (α) was estimated by averaging the black- and white sky surface albedo of shortwave radiation, following Duveiller et al.^2^.

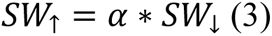

The shortwave reflected radiation was downscaled from 1° to 0.05°, with variations in downwelling shortwave radiation assumed to be homogenous across neighboring regions. Longwave emitted radiation at clear-sky conditions (LW^∗^) was calculated by using the Stefan-Boltzmann law:

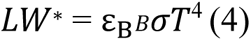

with the mean LST (T) from MODIS^6^, the Stefan-Boltzmann constant 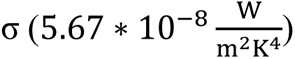 and the broadband emissivity *ε*_*B*_. Here, *ε*_*B*_ is estimated from the emissivity at 8400-8700 nm (*ε*_29_), 10780-11280 nm (*ε*_31_), and 11770-12270 nm (*ε*_32_) from MODIS MYD11C3^6^ by using the following formula^56^:

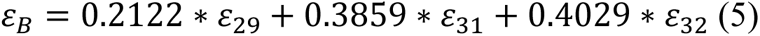

Changes in latent heat flux, longwave emitted radiation, and reflected shortwave radiation were calculated for each year combination by subtraction. As with LST, changes in the three components of the surface energy balance over a given period were conceptualized as the sum of the response due to ΔFC (ΔLE_fcc_, 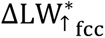, and 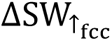) and the natural climate variability (ΔLE_res_, 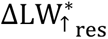, and 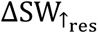).

Applying the same method as described above for ΔT, the residual changes in the components of the surface energy balance were estimated from adjacent stable forests and subtracted from the total response. This allowed us to estimate changes in the components of Earth’s surface energy balance due to Δ*FC*:

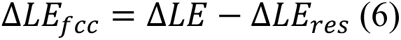

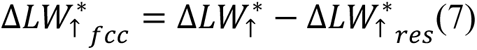

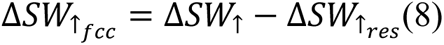

All estimated components of the surface energy balance represent all-sky conditions, except for the LST used for calculating 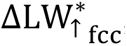, which was measured at clear-sky conditions. To correct for this, we multiplied 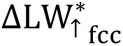 by the cloud correction factor (*CCF*), using a method modified from Duveiller et al.^2^. The cloud correction factor was calculated based on the all- and clear-sky (*) longwave emitted radiation from Kato et al.^55^ for the focal period. The *CCF* was downscaled from 1° to 0.05°:

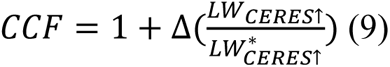

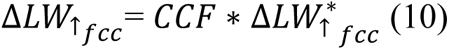

In total, the change in the Earth’s surface energy balance due to Δ*FC* during a certain period can be estimated as:

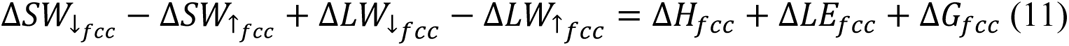

with changes in 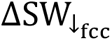 (downwelling shortwave radiation), 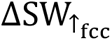 (shortwave reflected radiation), 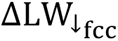 (downwelling longwave radiation), 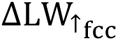 (longwave emitted radiation), ΔH_fcc_(sensible heat flux), ΔLE_fcc_(latent heat flux), and ΔG_fcc_ (ground heat fluxes) due to ΔFC. Following Duveiller et al.^2^, we assumed that changes in forest cover are too small to influence local cloud regimes. As such, 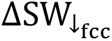 and 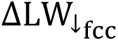 were set to zero.

With the previously calculated ΔLE_fcc_, 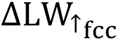, and 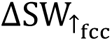, we were able to estimate the alterations in the combined ground and sensible heat fluxes due to ΔFC (Δ(H + G)_fcc_) by calculating the full energy balance at 0.05°. Based on^2^, we used the following formula to calculate Δ(H + G)_fcc_:

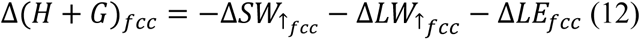

Finally, to convert the units of the surface energy balance components, all components were divided by 4*σT*^3^, where *T* is the mean LST (°K) of the earliest year of the focal period (i.e., 2003, 2004, or 2005)^2^. We scaled the changes in mean LST due to the components of the Earth’s surface energy balance so that negative (positive) values indicate a cooling (warming) effect due to energy loss (absorption) by the Earth’s surface.

To show the relative contribution of the different components of the Earth’s surface energy balance as a function of latitude, ΔT due to alterations in particular energy fluxes were upscaled to 4° resolution and then averaged per 4° latitudinal intervals (Fig. S1). The relative contributions were estimated per 4°-latitudinal interval by dividing the absolute value of a particular surface energy balance component by the sum of the absolute values of all components of the surface energy balance. Latitudinal intervals with less than three data points were excluded to limit the uncertainty of the estimates.

### Multivariate linear regression analysis

We used multivariate linear regression analysis to quantify relationships between Δ*FC* and ΔT and to quantify radiative and heat fluxes due to Δ*FC* (Δ*T*_*fcc*_), while controlling for other important environmental variables (i.e., background climate^27,28^, topography^29^, and soil type^30^). The Earth surface energy fluxes were not used for modeling the effects of Δ*FC* on ΔT since these fluxes do not not respond independently from one another. Furthermore, we assumed that the absolute effect of forest gains and losses on Δ*T*_*fcc*_ were equivalent and linear, following previous studies^1,5^.

#### Variable selection

We began with climatic^27,28^, topographic^29^, and ecological variables^30^ chosen a priori based on their expected relevance in explaining ΔT_fcc_ (Table S1). Climatic variables came from the climatologies at high resolution for the Earth’s land surface areas (CHELSA) data^27,28^. Yearly CHELSA data were averaged over the 2003-2012 period. Additionally, we used the snow water equivalent data of the European Space Agency (ESA), which is available for the period 1980-2018^31^. We took the annual sum of the 2-daily measurements to obtain the total snowfall per year.

ΔT_fcc_ and ΔFC were linked to the selected environmental parameters in Google Earth Engine based on their coordinates^51^. Based on the WWF terrestrial ecoregion classification^57^, the data were grouped into 4 regions: boreal (Boreal Forests/Taiga, and Tundra), temperate (Temperate Broadleaf & Mixed Forests, Temperate Conifer Forests, Temperate Grasslands, and Savannas & Shrublands), arid (Mediterranean Forests, Woodlands & Scrub, and Deserts & Xeric Shrublands), and tropical (Tropical & Subtropical Moist Broadleaf Forests, Tropical & Subtropical Dry Broadleaf Forests, Tropical & Subtropical Coniferous Forests, and Tropical & Subtropical Grasslands, Savannas & Shrubland). The biomes ‘Flooded Grasslands & Savannas’, ‘Montane Grasslands & Shrublands’, and ‘Mangroves’ were excluded from the analysis due to a low number of data points and high environmental distinctness (e.g., flooding frequency, elevation, etc.) compared to the other ecoregions. Further analysis was performed in R^58^.

To avoid multicollinearity, we then performed variable selection on the whole dataset. After scaling the variables by subtracting the mean and dividing by the standard deviation, we performed hierarchical clustering with the R package *ClustOfVar*^59^. Tree cutting was done by using an optimal number of clusters inferred from Principal Component analysis to remove correlated variables. The optimal number of clusters was estimated by taking the minimum number of eigenvectors needed to explain ≥95% of the cumulative proportion of the explained variation. Then, we checked the variance inflation factor (VIF) of the remaining variables and removed variables with VIF ≥ 4.

Multivariate linear models were run separately for the boreal, temperate, arid, and tropical regions. To further avoid multicollinearity in these regional sub-models, variables with VIF ≥ 4 in any of the regional models (excluding interaction terms) were removed from all sub-models. Since snow is an important determinant of the effect of ΔFC on the local climate in colder regions due to its effect on albedo^47,60^, we always included the snow water equivalent as an independent variable in models for the boreal and temperate regions.

#### Linear regression analysis

The final models of changes in land surface temperature included the selected variables (*X*), ΔFC, and the interactions between *X* and ΔFC as predictors. For the period 2003-2012, our statistical models consisted of the following independent variables *X* selected based on clustering and VIF (see above): mean annual temperature, mean precipitation of driest quarter ^27,28^, topographic surface roughness^29^, soil sand content^30^, and the sum of snow water equivalent^31^.

We normalized the independent variables *X* by subtracting the mean and dividing by the standard deviation. The dependent variables were the ΔFC-induced changes in the day, mean, and night LST, and changes in mean surface temperatures caused by the different components of the surface energy balance (i.e., shortwave reflected radiation, longwave emitted radiation, latent heat flux, and combined ground and sensible heat fluxes).

The intercept *a* of the models was forced through zero since it can be expected that ΔT_fcc_ equals zero when ΔFC ≅ 0 (equation 13)^1^. With a model structure that doesn’t consider the interaction between the precipitation variable(s) and snow water equivalent, it is predicted that a decrease in snow results in an increase in surface warming in boreal and temperate regions. However, this contrasts with previous studies showing the opposite response^3–5^. Including this interaction term in both biomes revealed that forests warm the surface less with decreased snow cover since accounting for the changes in the type of precipitation. Furthermore, across models predicting changes in land surface temperatures and surface energy fluxes between 2003 and 2012, including the interaction term always increased model R-squared, with a range of 0.2 −14% and 0.5 – 5% improvement in the boreal and temperate region, respectively. Accordingly, we draw our inference from models including the interaction term between the precipitation variable(s) and snow water equivalent in the boreal and temperate regions (equation 14).

Arid and tropical biome:

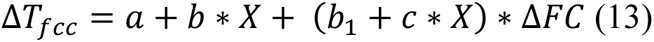

Boreal and temperate biome:

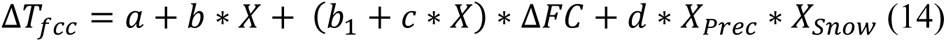

with a = 0, b = [b_2_ ⋮ b_n+1_], X = [X_1_ ⋮ X_n_] (with X_Prec_ and X_Snow_ representing the precipitation variables and the snow water equivalent, respectively), c = [c_1_ ⋮ c_n_], and *d* the coefficient of the interaction term between the precipitation and snow variables.

We tested the multivariate linear models for homoscedasticity using the Breush-Pagan test of the R package *lmtest*^61^. In case heteroscedasticity occurred (P-value < 0.05), the standard errors and the statistical significance of the predictors (F- and P-values) were corrected by using the variance-covariance heteroscedasticity consistent type 3 (HC3) of the R package *sandwich*^62,63^.

All multivariate linear models were statistically significant (P-value < 0.001) and explained, on average, 21 % (range: 3 - 35 %) of the variation in local surface temperature changes and 20 % (range: 5 - 47 %) of the variation in radiative and heat flux changes (Table S2).

#### Spatial autocorrelation

While spatial autocorrelation between residuals does not bias coefficient estimates in linear models, they can affect the results of statistical significance tests^64,65^. To test whether our model residuals showed problematic spatial autocorrelation, we used the Moran’s I test from the R package *moranfast* (https://github.com/mcooper/moranfast). Spatial autocorrelation was minor in most cases, particularly in the tropical biome (Fig. S3).

Directly accounting for spatial autocorrelation is computationally prohibitive with large datasets because the associated pairwise distance matrices are immense. Accordingly, to test whether the output of the models qualitatively differs if accounting for spatial autocorrelation, we compared the standard linear model output of 2003-2012 with the output of a spatial lag and error model based on a subset of 15,000 data points per biome. We used the R package *spdep* for spatial regression analysis^66^ (Fig. S3; Fig. S4). For comparison, a multivariate linear model was also generated on the biome-level data subsets. Model standard errors of the coefficients and their significance values were corrected with the occurrence of heteroscedasticity using HC3^62,63^.

To assess the potential influence of spatial autocorrelation on our findings, we quantified the percentage of model variables with the same sign per biome across all variables of interests. We showed that for the period 2003-2012 model coefficients describing the temperature response to ΔFC and its interaction with environmental variables were most similar in terms of sign in the temperate region (81%), followed by the boreal (78%), arid (73%), and tropical regions (67%).

#### Model validation

We tested the performance of the biome-level linear models for the period 2003-2012 by predicting ΔT_fcc_ as a response to ΔFC using new input data. The new input data corresponded to 15-year combinations with different time lengths, specifically every year between 2004 and 2012 with 2003 as a baseline, and the nine different year combinations between 2003-2005 and 2010-2012. Variables were normalized using means and standard deviations of the data from the 2003-2012 model. To assess the discrepancy in observed and predicted ΔT_fcc_, we performed linear regression of the observed versus predicted ΔT_fcc_ per climate region (Fig. S2).

#### The background climate

We used the linear models from 2003-2012 to assess the role of background climate in determining the effect of forest cover change on local temperatures (Fig. 2). To extract the effect of each climate variable (i.e., mean annual temperature, mean precipitation of driest quarter, and sum of snow water equivalent) on the ΔFC-ΔT_fcc_ relationship, biome-level model partial residuals were extracted for the interaction term and plotted as a function of ΔFC^67^. The predicted values of the mean values of the predictors were added to the partial residuals. Thereafter, we performed global-level linear regression analysis per quantile category of precipitation and temperature (<0.1, 0.1-0.9, and >0.9). Additionally, we distinguished between regions with (>0.01 mm) and without snowfall (<0.01 mm). We then determined the significance values of the interaction between the climate variable of interest and ΔFC (while accounting for heteroscedasticity; see above) based on a global-level model.

### The sensitivity of local surface temperatures to forest cover change

#### Spatial and temporal analysis

To calculate the slope between ΔT_fcc_and ΔFC (i.e., ΔT/ΔFC) for each pixel, we multiplied the interaction term coefficients from our multivariate models with the pixel’s normalized variables

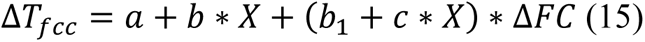

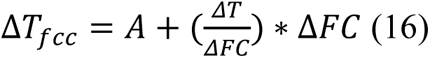

where *A* = *a* + *b* ∗ *X* and 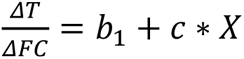.

We conducted the analysis using the models based on the period from 2003 to 2012 as longer periods result in more robust estimates of temperature responses to forest cover changes^1^. ^1^Lastly, we validated the model with data from different year combinations (Fig. S2).

The values of the variables X were drawn from the same global gridded data products used in the models. For CHELSA^27,28^ and ESA^31^ data, we used yearly averages between the years 1980 and 2018. The final analysis covered the years 1988-2016 as observations of snow water equivalent decreased drastically in quantity and quality this period due to a change in the satellite type (Fig. S10a). The years 2017 (extremely wet year) and 2018 (dry year) were omitted from the final analysis due to climate anomalies (i.e., outliers in the linear models predicting the proportion of land cooling under forests as a function of time).

With our approach, we were able to quantify how the changing background climate influences the impact of ΔFC on LST across these 28 years. The pixel-level variables were normalized as in the multivariate linear models. ΔT/ΔFC was predicted for tropical, temperate, boreal, and arid regions that contained forest (defined as pixels with > 10% canopy cover) in 2019^7,18^. The pixel-level quantification of ΔT/ΔFC was done in Google Earth Engine^51^. The change in ΔT/ΔFC over the past three decades was calculated as the difference between ΔT/ΔFC in 2016 and 1988 (Fig. 3; Fig. S13). Mapping was done in R^58^, whereby values outside the 80% confidence interval of the mean were set to the 0.1 or 0.9 quantiles.

#### Inter-annual variability

Inter-annual climate variability potentially affects estimating temperature responses to vegetation changes. To test this, our approach was two-fold (based on Alkama and Cescatti^1^). Firstly, ΔT/ΔFC was estimated between the fixed year 2003 and the years from 2004 until 2012 (Fig. S8). This was done by performing multivariate linear regression and calculating ΔT/ΔFC at locations with significant ΔFC in that particular year combination (using the method described above). The climatic variables used in the linear models for the different year combinations were the averages of the period 2003-2012. Secondly, ΔT/ΔFC was estimated based on the data corresponding to the nine different year combinations between 2003-2005 and 2010-2012 (Fig. S9). The averages and 95% confidence interval of ΔT/ΔFC per latitudinal zone (4°) were plotted for the year combinations between 2003-2005 and 2010-2012. To conclude, both analyses showed that the 2003-2012 period is well-suited for analyzing temperature responses to vegetation changes as it does not include highly anomalous years^1^.

#### The background climate

We used the multivariate linear models including several climate variables to estimate their contribution to ΔT/ΔFC between 1988 and 2016. To assess the impact of each variable, we ran simulations for the years 1988 to 2018 changing one variable at a time, while keeping the other uncorrelated variables constant at their mean values over the same period.

From these maps, we estimated the annual average ΔT/ΔFC (Fig. S11) and the proportion of negative ΔT/ΔFC under two scenarios: full climate change and change in only one climate variable at a time (Fig. 4). Additionally, we examined the role of each climate variables in determining ΔT/ΔFC by estimating the average change in mean ΔT/ΔFC per 4°-latitudinal intervals (Fig. S14).

Then, we calculated the percentage change of the global surface area where forests have a cooling effect (ΔA%) as follows:

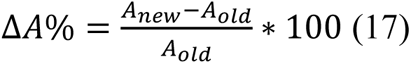

with *A*_*old*_and *A*_*new*_representing the surface area where forests have a cooling effect in 1988 and 2016, respectively. Using the same equation, we also estimated the percentage change in the annual average ΔT/ΔFC between 1988 and 2016.

#### Large-scale atmospheric circulation effects

To investigate the potential impact of atmospheric circulations on the local climatic effects of forests, we analyzed the relationship between land surface temperature sensitivities (ΔT/ΔFC) and changes in surface albedo. Using MODIS MCD43C3 data on surface albedo between 2003 and 2012^55^, we assessed the sensitivities through both raw data and model predictions to explore the relationship between surface temperature sensitivities and the alterations in surface albedo resulting from changes in forest cover.

Our results show a consistent slope in the relationship between surface albedo and surface temperature sensitivity during night-time across different latitudinal intervals and a comparatively weaker relationship for mean surface temperatures (Fig. S16). The consistent slope across different latitudes suggests that the confounding factors influencing this link are independent of latitude. Considering that circulation-induced changes are unlikely to act uniformly across latitudes^55^ we conclude that the effects of atmospheric circulations on the relationship between surface albedo and surface temperature sensitivity are negligible. Notably, during day-time, we observed a difference in the slope direction between tropical regions and higher latitudinal areas. This disparity underscores the need for further investigation to identify the specific confounding factors at play to develop a comprehensive understanding of the intricate complexities involved in this relationship.

#### Uncertainty of sensitivities

The uncertainty of the calculated ΔT/ΔFC was calculated by applying error propagation to the standard errors of the coefficients of the linear models and the scaled values of *X* (Fig. S12). *X* was assumed to be a constant since there were no standard errors available for these variables.

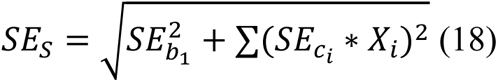

where SE_S_represents the standard error of the estimated pixel-level ΔT/ΔFC, 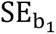 represents the standard error of the coefficient of ΔFC in the linear model (the standard error of the coefficient of the interaction terms in the linear model, and X_i_represents the values of the scaled variables. The mean of the standard errors was taken for the period 1988-2016 (Fig. S12).

## Acknowledgments

This work was supported by grants to T.W.C. from DOB Ecology and the Bernina Foundation. W.W.M.V. was supported by funding from the Ecosystem Management Group at ETH Zürich, and the FWO and F.R.S.-FNRS under the Excellence of Science program (EOS O.0026.22) at Ghent University. G.R.S. acknowledges funding from DOB Ecology, Bernina Initiative and from the Marc R. Benioff Revocable Trust, which, in collaboration with the World Economic Forum, made this work possible. C.M.Z and L.M. were funded by the Ambizione grant PZ00P3_193646. D.S.M. was supported by the Ambizione grant from the Swiss National Science Foundation to DSM (#PZ00P3_193612).

## Author Contributions

The study was designed by W.W.M.V., G.R.S., L.M., T.L., and T.W.C. Data analysis and interpretation, and manuscript writing was carried out by W.W.M.V. with support from L.M., G.R.S., T.L, C.M.Z., D.S.M., S.S., and T.W.C. G.R.S. and L.M. contributed equally to the paper and share the second authorship. All co-authors read and approved the manuscript.

## Competing interest declaration

The authors declare no competing interests.

## Materials & Correspondence

Correspondence should be addressed to W.W.M.V., G.R.S., and L.M. Requests for materials and code should be addressed to W.W.M.V.

## Code and data availability

All data and code will be shared on the online repository Zenodo.

## Extended Figures

### The Earth’s surface energy balance and forest cover changes

**Fig. S1:**
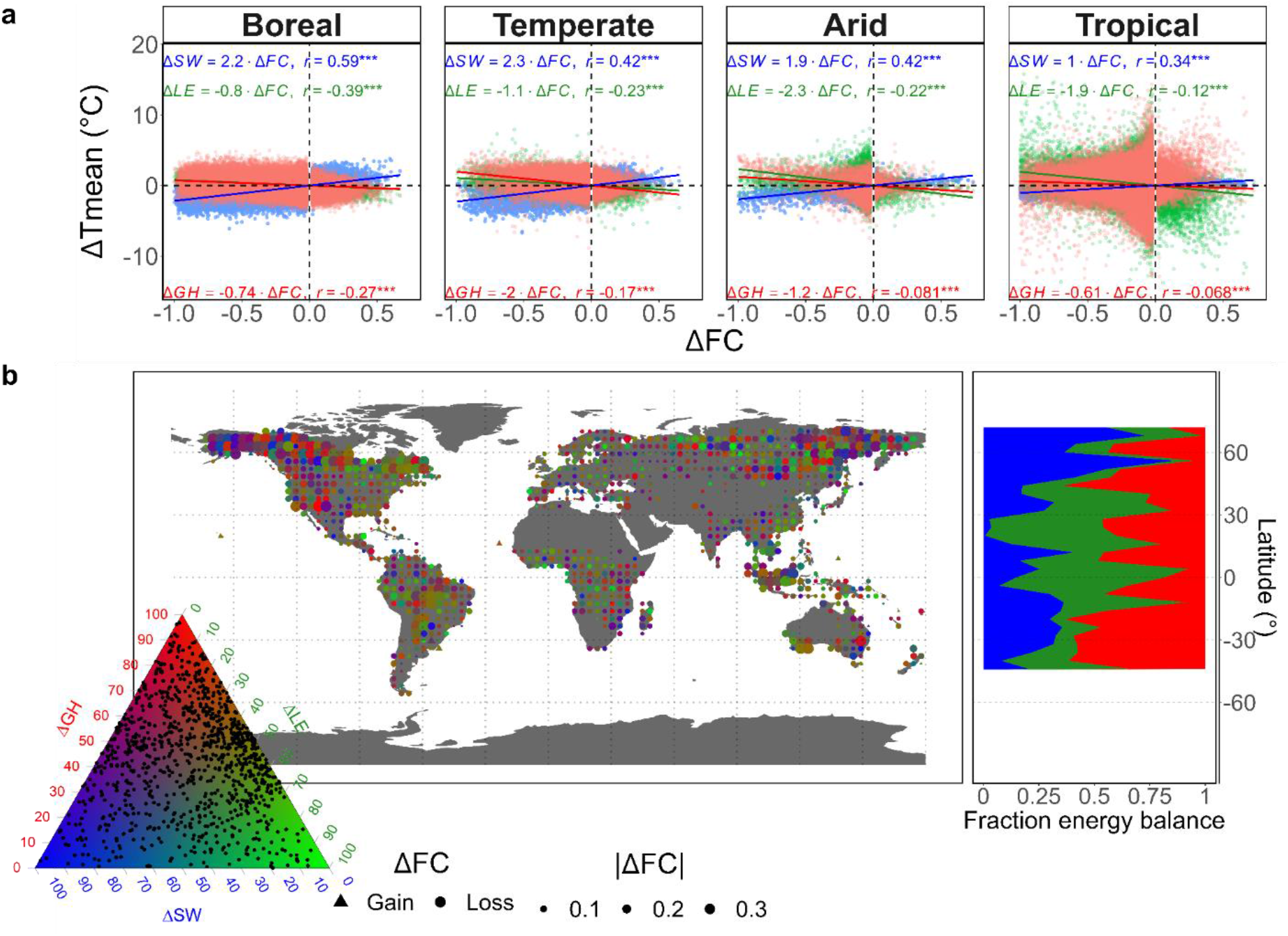
The impact of forest cover changes on the Earth’s surface energy balance for the period 2003-2012. **a,** The effect of forest cover change (Δ*FC*) on changes in mean surface temperature (Δ*Tmean*) due to alterations in shortwave reflected radiation (Δ*SW*), latent heat fluxes (Δ*LE*), and combined ground and sensible heat fluxes (Δ*GH*) for the boreal (n=195,892), temperate (n=158,851), arid, (n=17,598) and tropical biome (n=278,587). The coefficients of univariate linear models (Δ*T* = 0 + *β* ∗ Δ*FC*), and the Pearson correlation coefficient (r) and significance are shown. **b,** The relative contribution of the three components of the Earth’s surface energy balance to Δ*Tmean* (left) (n=1,014). The direction and magnitude of the forest cover changes are indicated by the shape and size of the symbols, respectively. The legend shows the tricolor plot with the location of the data points shown. The latitudinal pattern of the fraction of the energy balance (right). The average relative contribution of the Earth’s surface energy balance to Δ*Tmean* are shown per 4° latitudinal interval (n=1,009). Latitudinal intervals with less than 3 data points were excluded.

### Multivariate linear regression

**Fig. S2:**
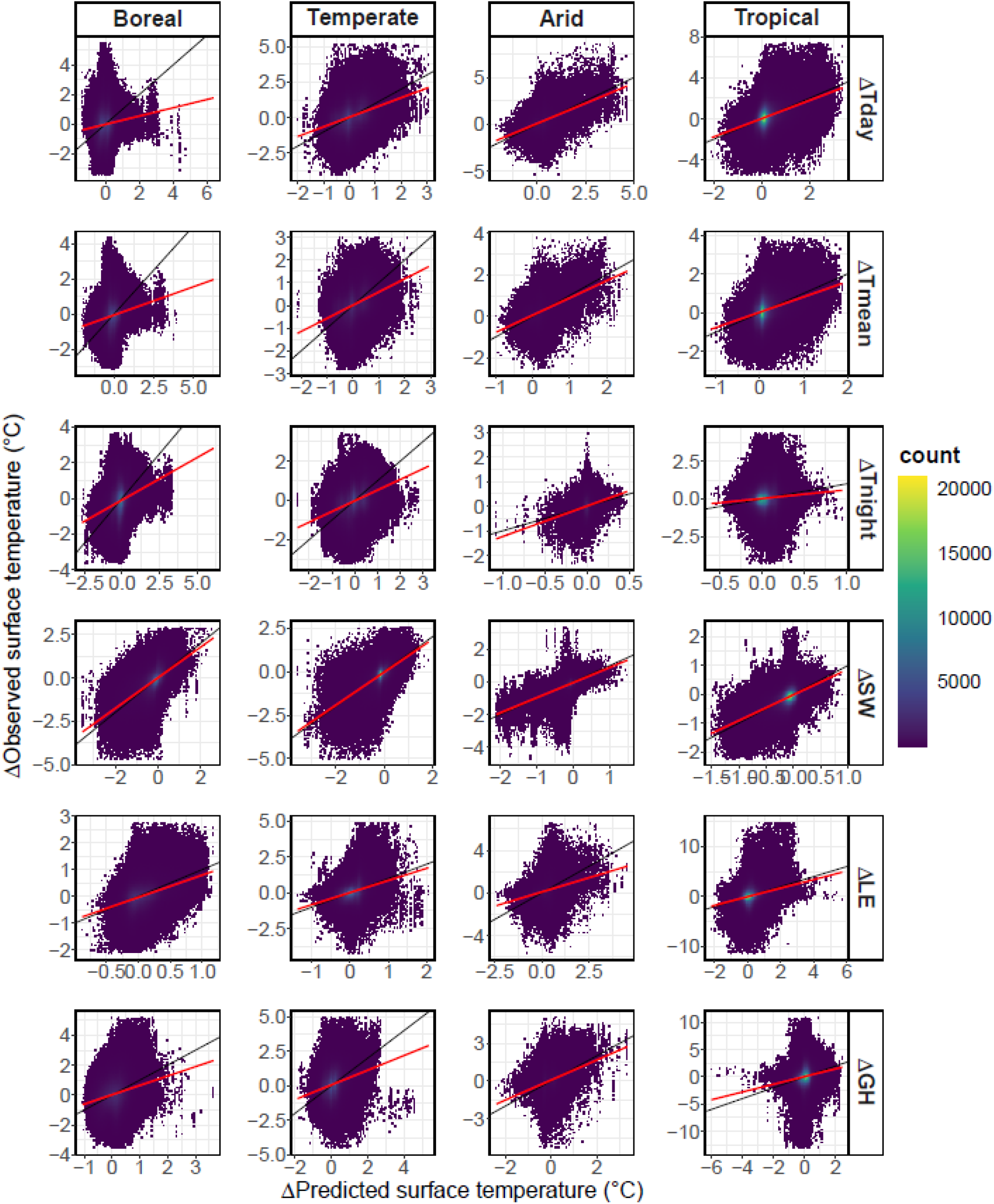
Validation of the models of the period 2003-2012. Linear regression was done on all data per biome and variable. Data not included in the interval defined by the lower and upper 0.00005 quantiles of observed and predicted surface temperatures were excluded per biome and variable for plotting purposes. The colour indicates the density of the points.

**Fig. S3:**
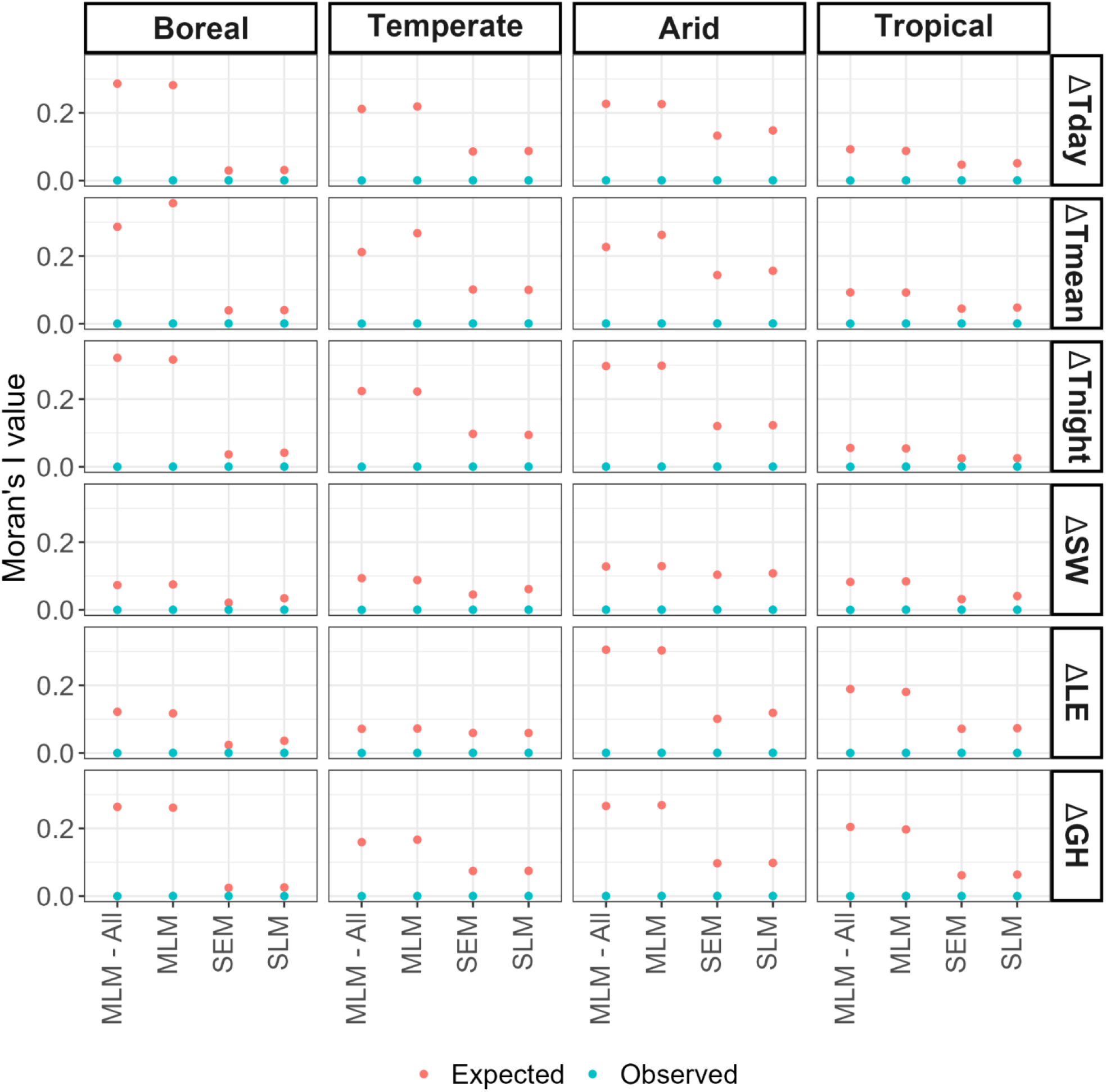
Moran’s I test for spatial models for the period 2003-2012. The expected and observed Moran’s I value for the multivariate linear models on the full dataset (MLM - All), and the multivariate linear model (MLM), spatial error model (SEM), and spatial lag model (SLM) on the subsetted data are shown. All tests were statistically significant (P-value < 0.001).

**Fig. S4:**
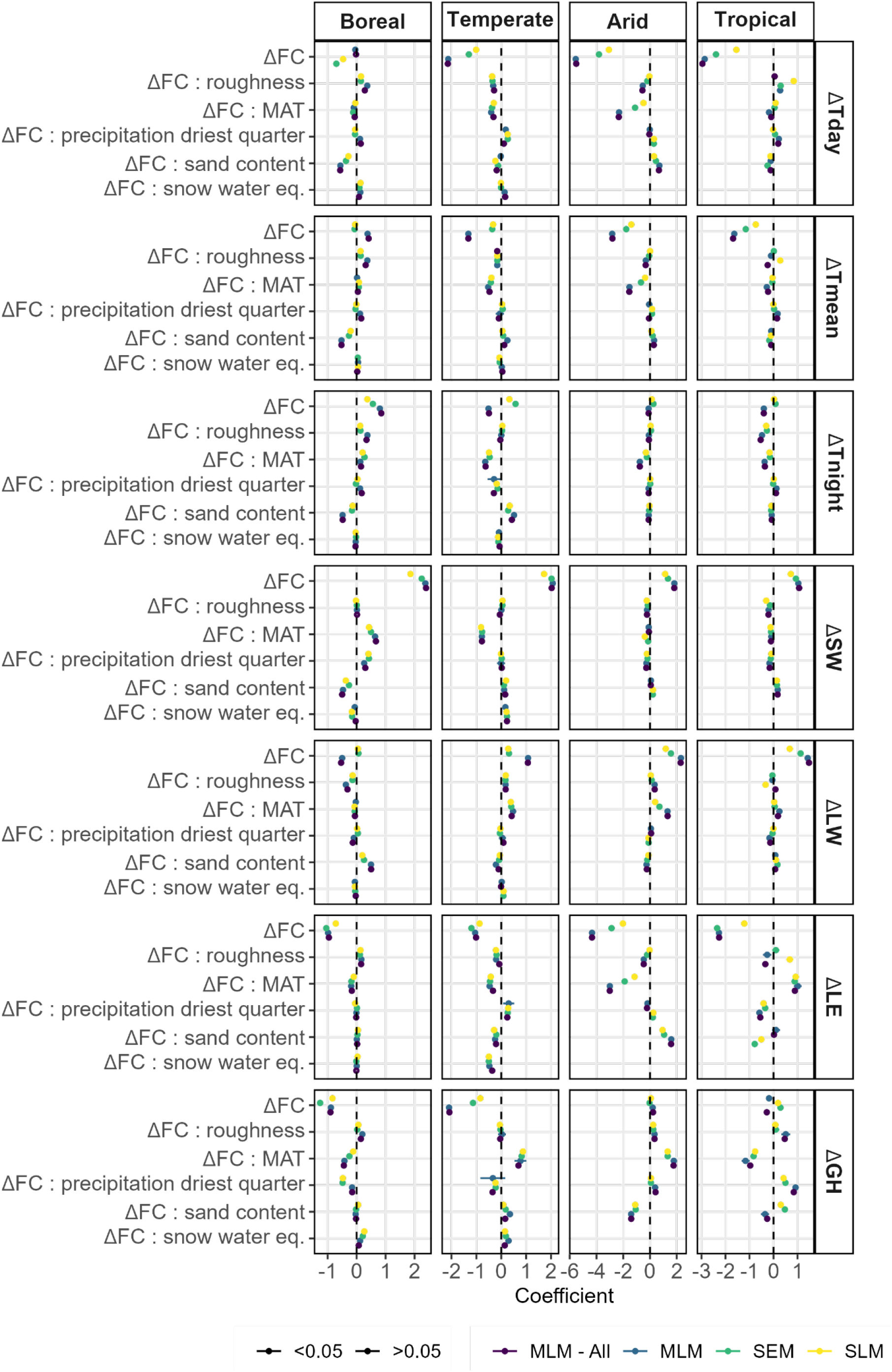
Validation of the effect of spatial autocorrelation on model. Model coefficients for the interaction terms and forest cover change are shown per biome and dependent variable for the period 2003-2012. Models included multivariate linear models based on all data (MLM - All) and on a subset of data (MLM), as well as spatial error (SEM) and lag models (SLM) on a subset of data. Data subsets were 15,000 randomly selected data points per biome. Points were filled in case model coefficients were significant in the model (heteroscedasticity was accounted for in multivariate linear models).

### The background climate

**Fig. S5:**
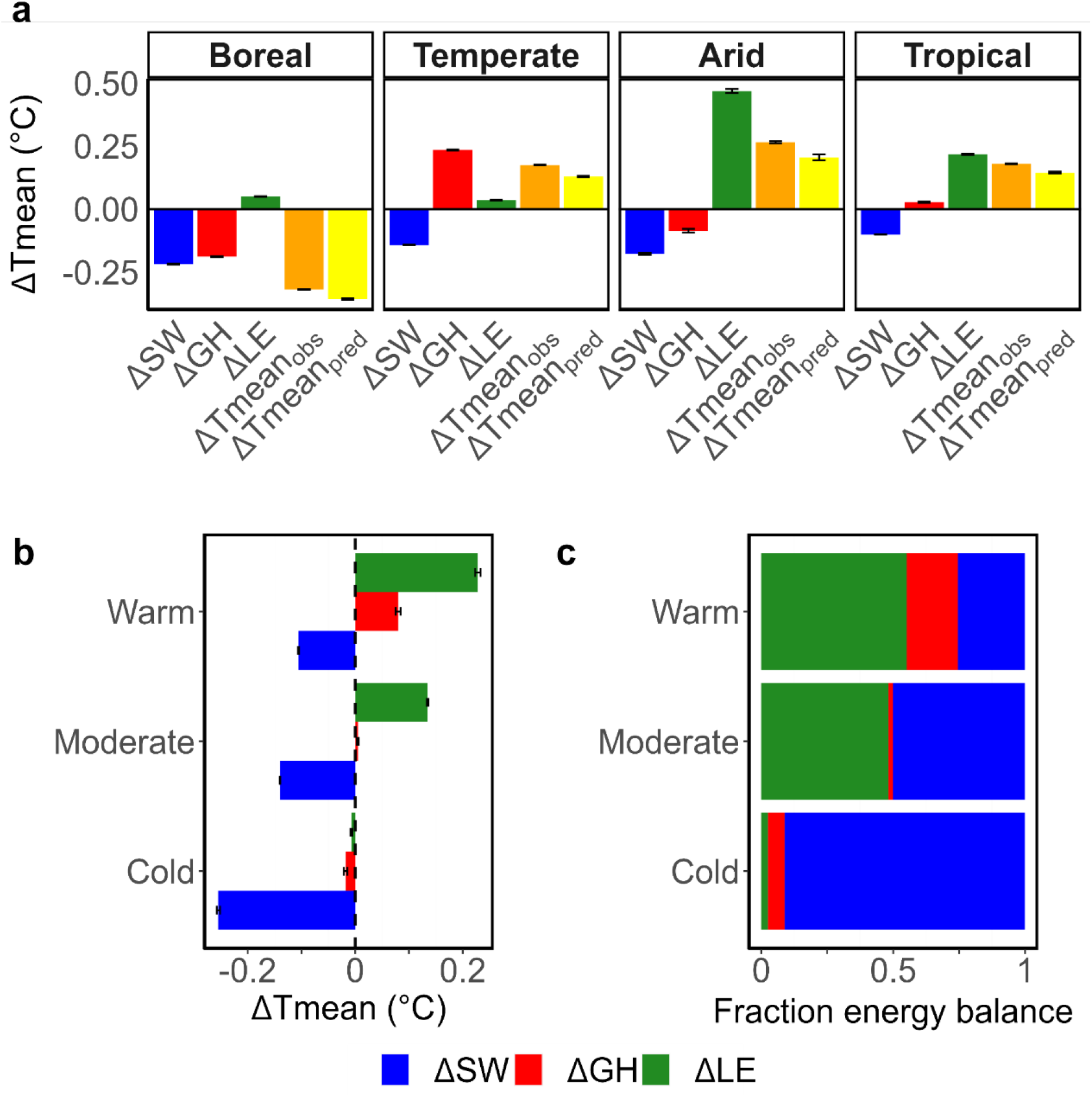
The influence of mean annual temperatures on the Earth’s surface energy balance for forest cover changes in 2003-2012. **a,** The averaged changes in mean surface temperatures due to alterations in the components of the Earth’s surface energy balance (shortwave reflected radiation: Δ*SW*; latent heat fluxes: Δ*LE*; combined ground and sensible heat fluxes: Δ*GH*) due to forest cover change between 2003 and 2012 for the boreal (n=195,892), temperate (n=158,851), arid, (n=17,598) and tropical biome (n=278,587). The average observed change in mean surface temperatures is shown (Δ*Tmeanobs*). The predicted mean surface temperature changes (Δ*Tmeanpred*) were calculated by summing Δ*SW*, Δ*GH*, and Δ*LE*. Error bars denote the standard errors. **b,** The average changes in mean surface temperature due to changes in Δ*SW*, Δ*LE*, and Δ*GH* with forest cover change per background temperature zone in 2003-2012. The cold (blue; n=65,093), moderate (grey; n=520,756), and warm (red; n=65,079) climates represent the 0-0.1, 0.1-0.9, and 0.9-1 quantile intervals of the average mean annual temperature of the period 2003-2012, respectively. Error bars denote the standard errors. **c,** Fraction energy balance contributing to the overall changes in mean surface temperatures with forest cover change per background temperature zone.

**Fig. S6:**
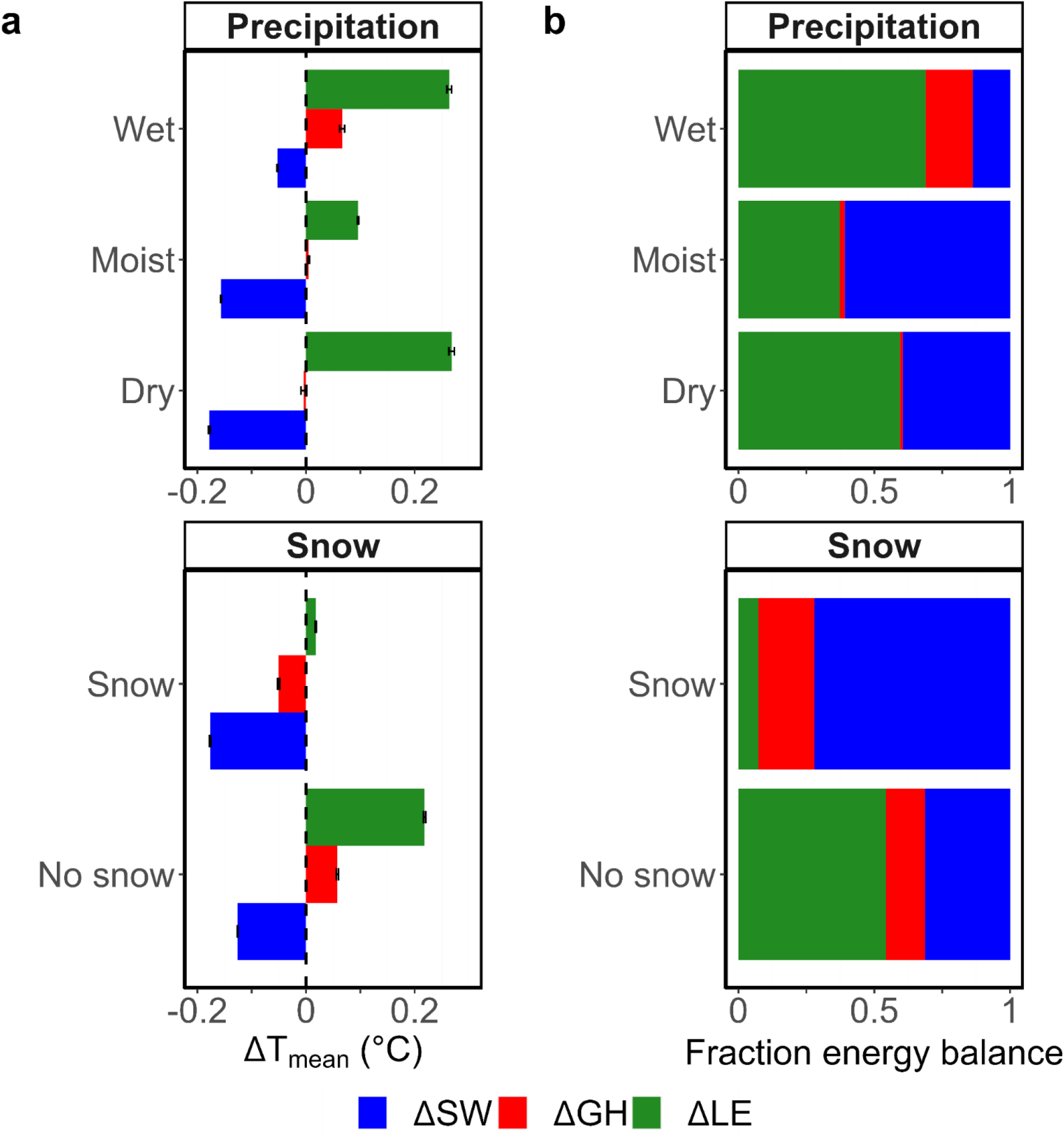
The background climate and the Earth’s surface energy balance for the period 2003-2012. **a,** Average changes in mean surface temperature due to changes in SW, GH, and LE per climate zone (top: precipitation of driest quarter; bottom: snow water equivalent) for the period 2003-2012. The dry (n=65,094), moist (n=520,741), and wet climates (n=65,093) represent the 0-0.1, 0.1-0.9, and 0.9-1 quantile intervals of the mean precipitation of the driest quarter of the period 2003-2012, respectively. Locations without (n=363,741) and with snow zones (n=287,187) were selected for the period 2003-2012. The error bars represent the standard errors. **b,** Fraction energy balance fraction contributing to the overall changes in mean surface temperatures per climate zone.

**Fig. S7:**
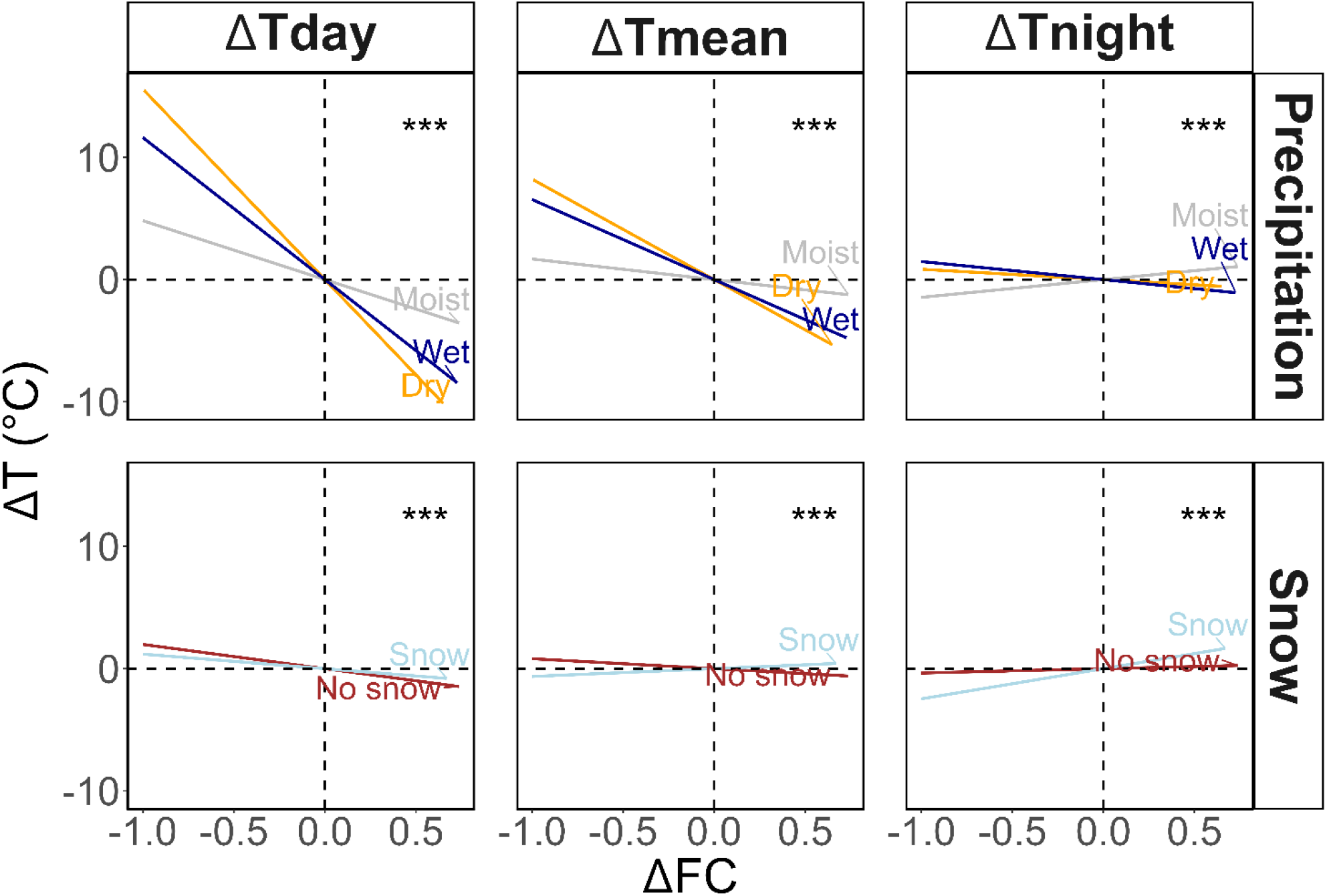
The influence of precipitation and snow on the surface temperature response to forest cover change. Biome-level model partial residuals in respect of the interaction between forest cover change and precipitation (mean precipitation of driest quarter) or snow (sum of snow water equivalent) for the models predicting day/mean/night surface temperature change (Δ*T*) as a function of forest cover change. The dry (n=65,094), moist (n=520,741), and wet climates (n=65,093) represent the 0-0.1, 0.1-0.9, and 0.9-1 quantile intervals of the mean precipitation of the driest quarter of the period 2003-2012, respectively. Locations without (n=363,741) and with snow zones (n=287,187) were selected for the period 2003-2012. Global-level linear regression was then performed per climate zone. The stars indicate the significance of the interaction between the climate variable and Δ*FC* from a global level combining all climate zones.

### Inter-annual variability

**Fig. S8:**
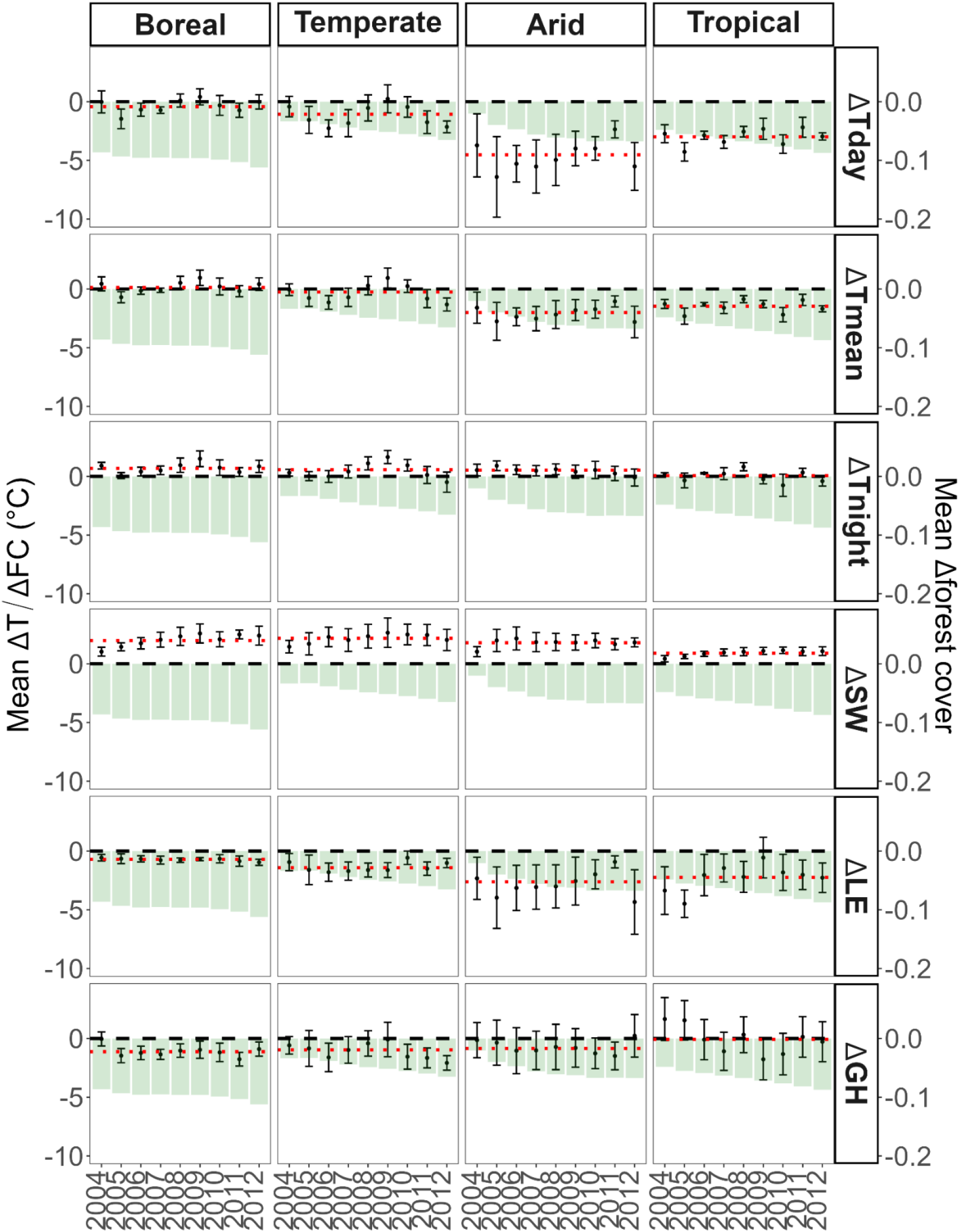
Inter-annual variability of the sensitivities of surface temperature and the components of the Earth’s surface energy balance to forest cover change. Sensitivity of day (Δ*Tday*), mean (Δ*Tmean*), and night surface (Δ*Tnight*) temperature, and the mean surface temperatures as a result of changes in shortwave reflected radiation (Δ*SW*), latent heat fluxes (Δ*LE*), and combined ground and sensible heat fluxes (Δ*GH*) to forest gain between a fixed year 2003 and the years 2004-2012 (shown as points). The error bars represent the standard deviation of the mean sensitivity. The red dotted line indicates the mean temperature sensitivity for the year combinations excluding the period 2003-2012. The second axis on the right shows the mean forest cover change for every period (shown as green bars). Positive (negative) values represent forest gain (loss).

**Fig. S9:**
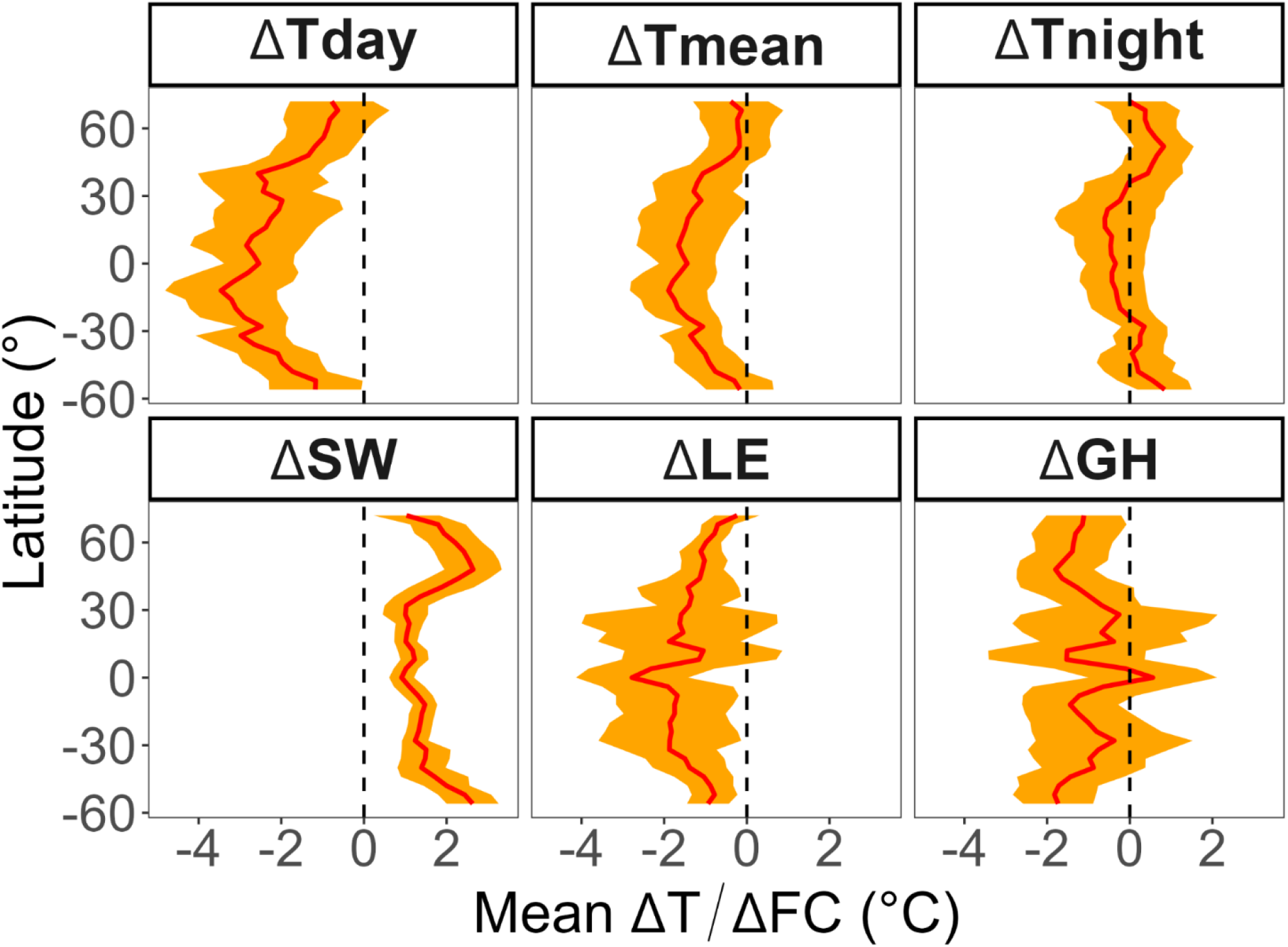
Latitudinal patterns of the sensitivities of surface temperature and the components of the Earth’s surface energy balance to forest cover change. Zonal averages (ΔT) of changes in day, mean, and night surface temperatures, and changes in mean surface temperatures due to changes in shortwave reflected radiation (Δ*SW*), latent heat fluxes (Δ*LE*), and combined ground and sensible heat fluxes (Δ*GH*) as a response to forest gains at 4 °C latitudinal resolution. The data are the sensitivities obtained from the nine different combinations between the years 2003-2005 and 2010-2012 at places with significant forest cover change. The standard deviation is given in orange.

### The sensitivity of local surface temperatures to forest cover change

**Fig. S10:**
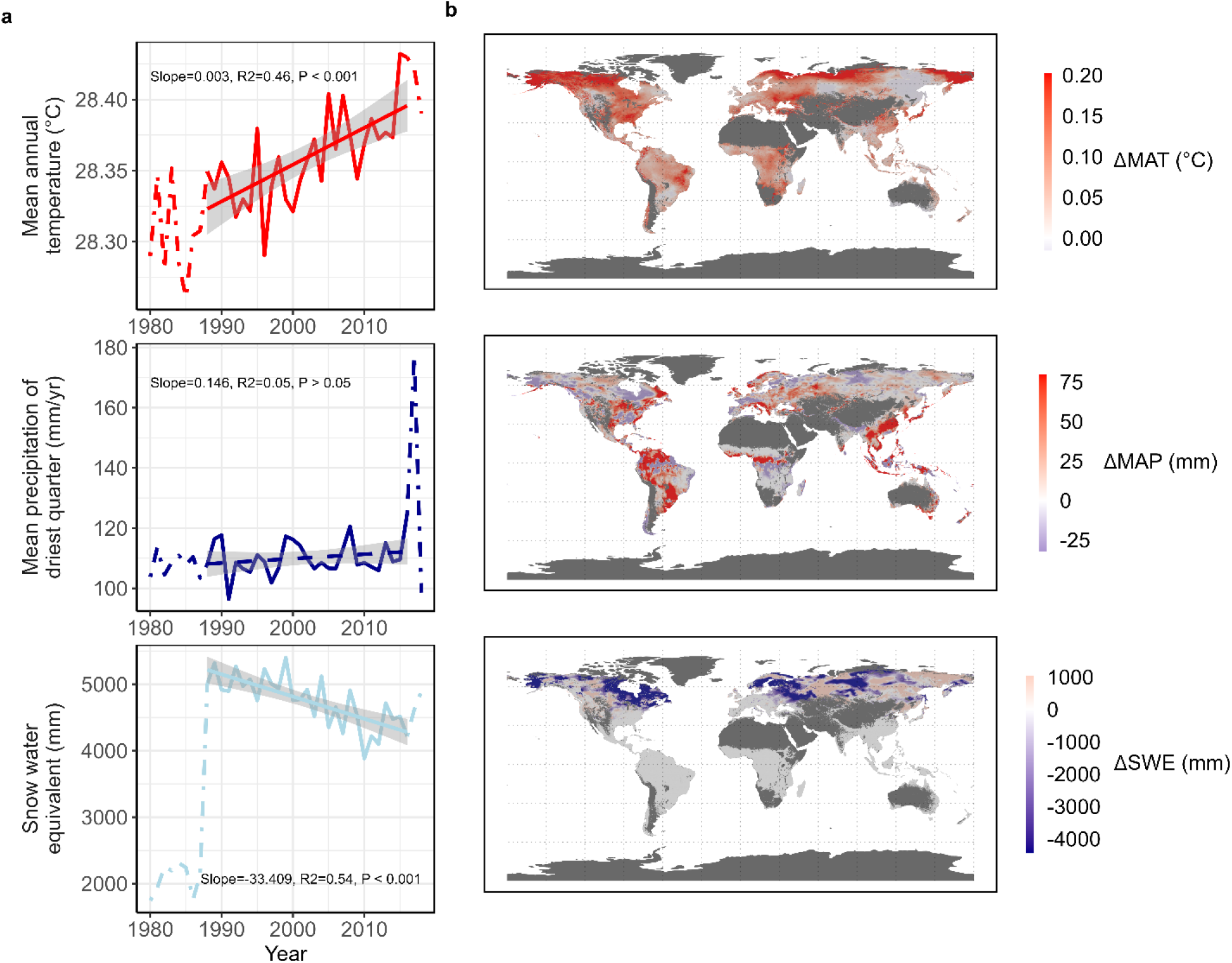
Change in the average mean annual temperature, mean precipitation of driest quarter, and sum of snow water equivalent over the past three decades. **a,** The global-level change in the average mean annual temperature, mean precipitation of driest quarter, and sum of snow water equivalent for the period 1980-2018. The dotted lines indicate the period excluded from the final analysis (i.e., 1980-1988 and 2016-2018). Linear regression analysis was performed for the period 1988-2016 (n=3,764,012). Solid lines represent regression lines with significant P-values (<0.05), while dashed lines denote non-significant regression lines (P-values >0.05). **b,** The spatial variation in climate change during the period 1988-2016 for the change in the mean annual temperature (ΔMAT), mean precipitation of the driest quarter (ΔMAP), and the sum of the snow water equivalent (ΔSWE) between 1988 and 2016. The 80% quantile interval was shown.

**Fig. S11:**
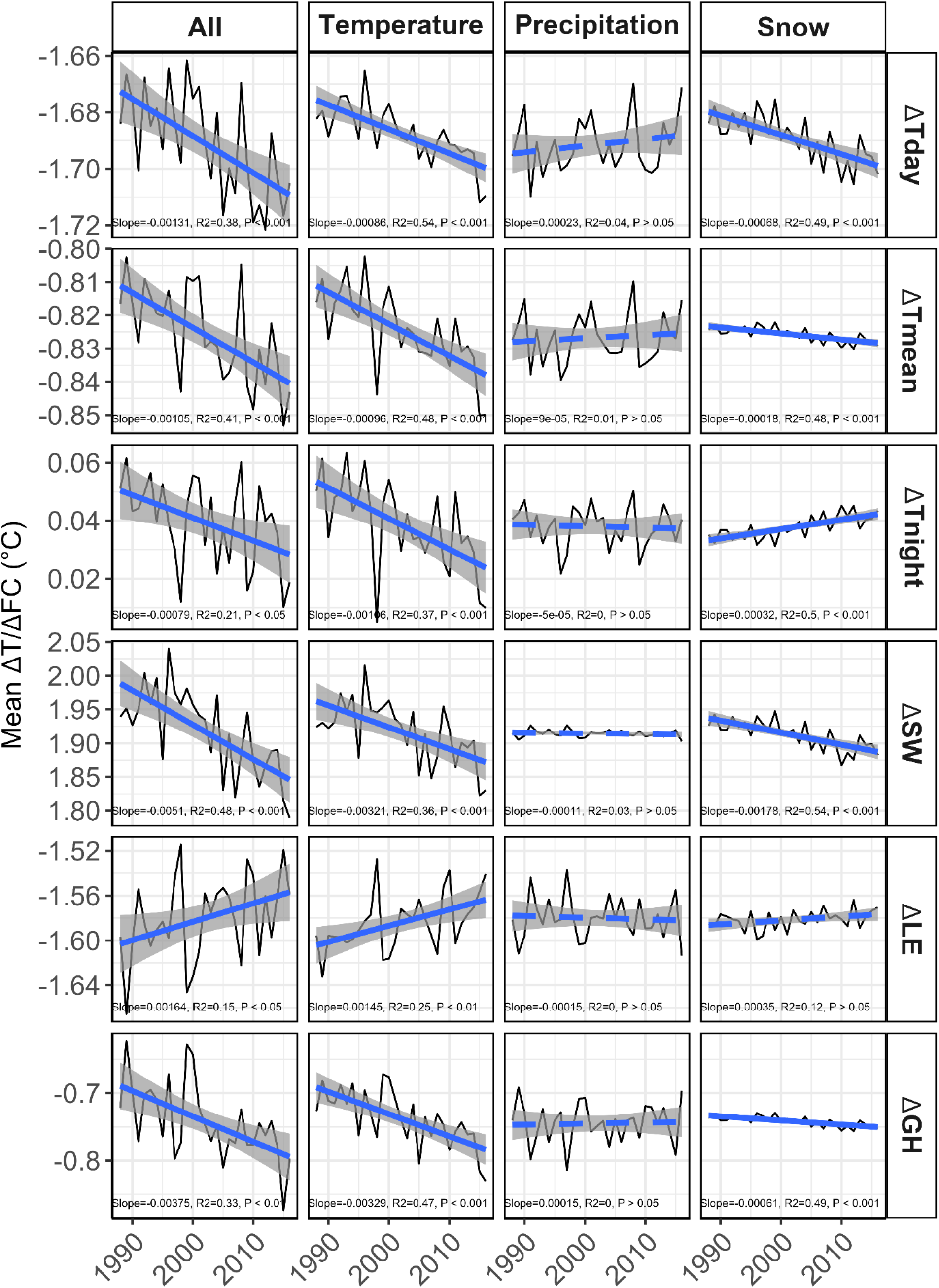
Global averages of land surface temperature and heat flux sensitivities to forest cover change (ΔT/ΔFC) for the period 1988-2016. The annual mean sensitivity of day (ΔTday), mean (ΔTmean), and night (ΔTnight), surface temperatures and the different components of the Earth’s surface energy balance (reflected shortwave radiation: ΔSW; latent heat fluxes: ΔLE; combined ground and sensible heat fluxes: ΔGH) were estimated. The overall change in sensitivities was estimated by using yearly temperature, snow, and precipitation variables (All) (n=3,764,012). To disentangle the influence of the different climate variables, one climate variable (Temperature, Precipitation, and Snow) was changed at a time while the other were kept constant using the average of the whole period. Significant regression lines (P-value < 0.05) are solid lines. Non-significant regression lines (P-value > 0.05) are dashed lines.

**Fig. S12:**
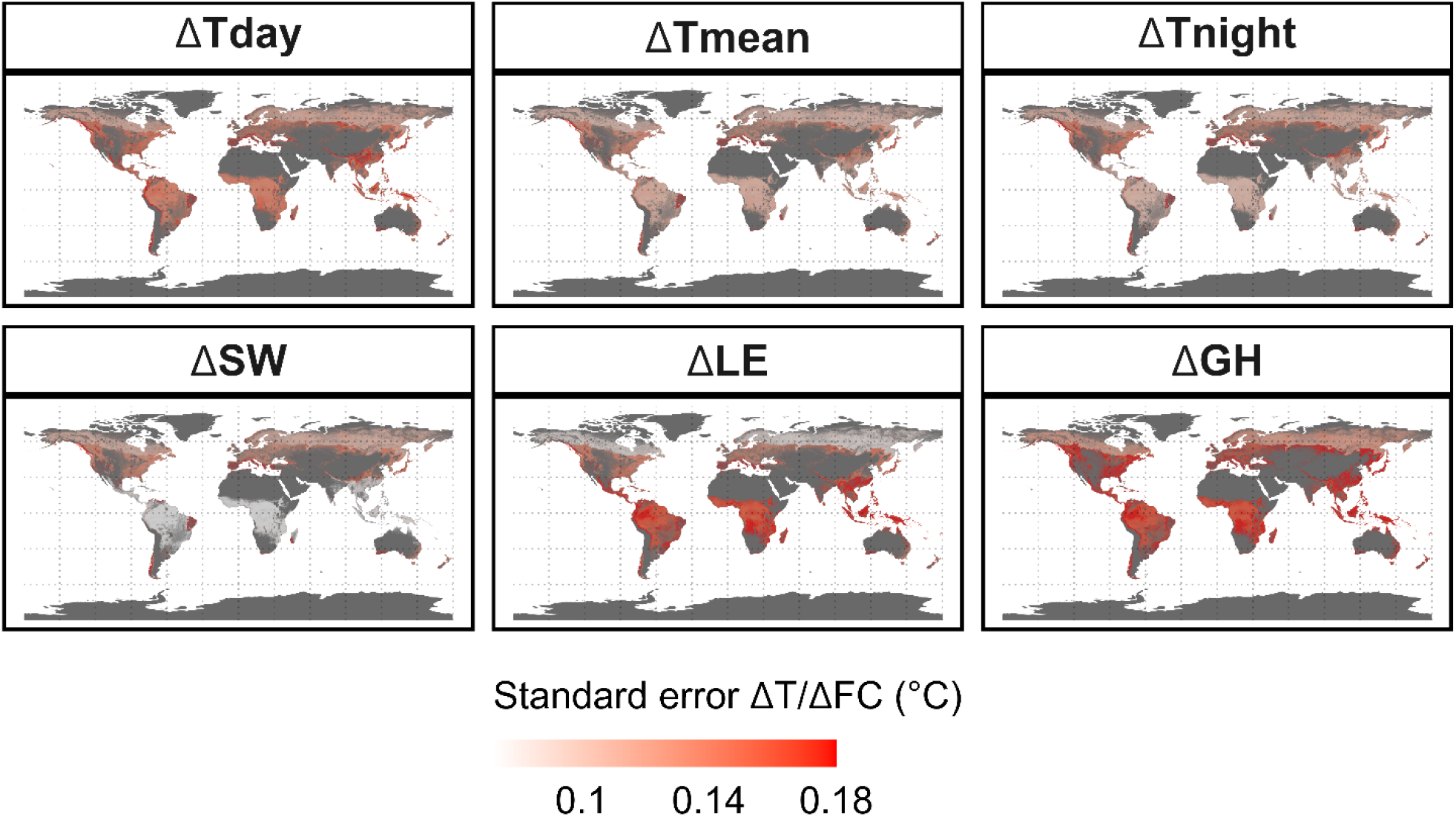
Standard errors of the sensitivity of day (ΔTday), mean (ΔTmean), and night temperature (ΔTnight), and shortwave reflected radiation (ΔSW), latent heat fluxes (ΔLE), and combined ground and sensible heat fluxes (ΔGH).

**Fig. S13:**
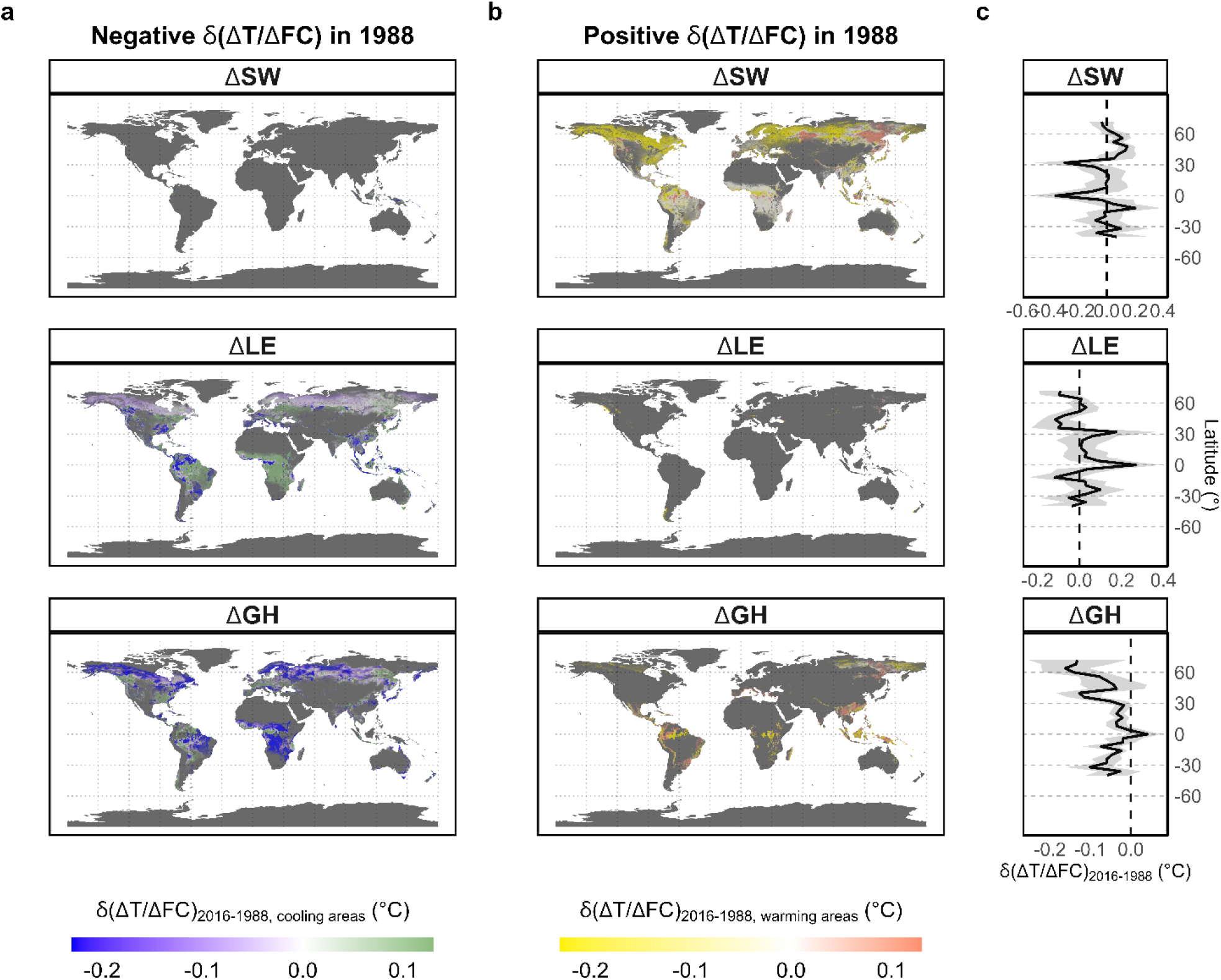
Sensitivities of Earth’s surface energy balance components to forest cover change between 1988 and 2016. **a,** Change in the sensitivity of the change in mean surface temperatures due to alterations in reflected shortwave radiation (Δ*SW*), latent heat fluxes (Δ*LE*), and combined ground and sensible heat fluxes (Δ*GH*) to forest cover change between 1988 and 2016 for pixels with negative sensitivities to forest cover change (i.e., areas cooling; n=2,719,786) in 1988. **b,** Change in the sensitivity of LE, SW, and GH to forest cover change between 1988 and 2016 for pixels with positive sensitivities to forest cover change (i.e., areas warming; n=1,044,226) in 1988. **c,** Latitudinal pattern of the difference in the sensitivity of LE, SW, and GH to forest cover change between 1988 and 2016 based on 4° latitudinal means. Latitudinal intervals with less than 3 data points were excluded. The 95% confidence interval is shown in grey.

**Fig. S14:**
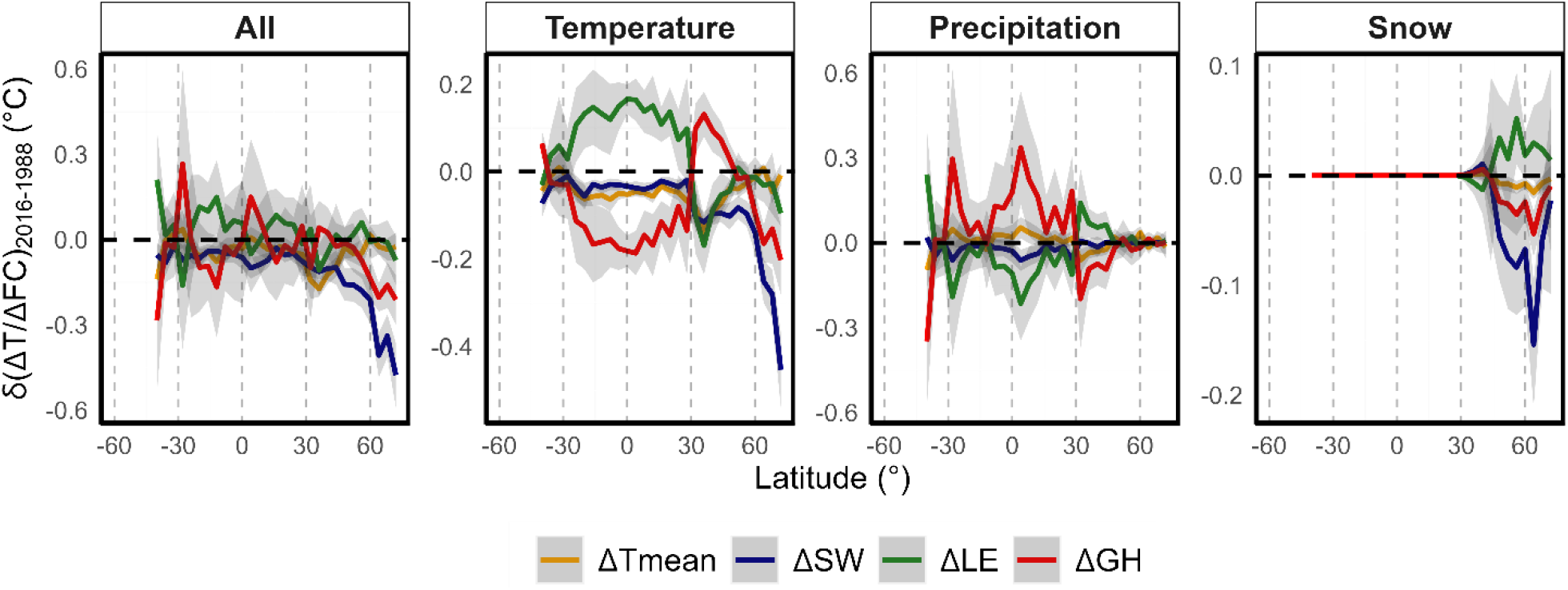
The latitudinal pattern of the response of the Earth’s surface energy balance to changes in the background climate between 1988 and 2016. The average change in the sensitivity of mean surface temperatures (Δ*Tmean*) and the contribution of the Earth’s surface energy balance components (shortwave reflected radiation: Δ*SW*; latent heat fluxes: Δ*LE*; combined ground and sensible heat fluxes: Δ*GH*) per 4°-latitudinal intervals (n=978). The effect of the changing background climate was shown for all climate variables (All), mean annual temperatures (Temperature), mean precipitation of driest quarter (Precipitation), and snow water equivalent (Snow). Latitudinal intervals with less than 3 data points were excluded.

**Fig. S15:**
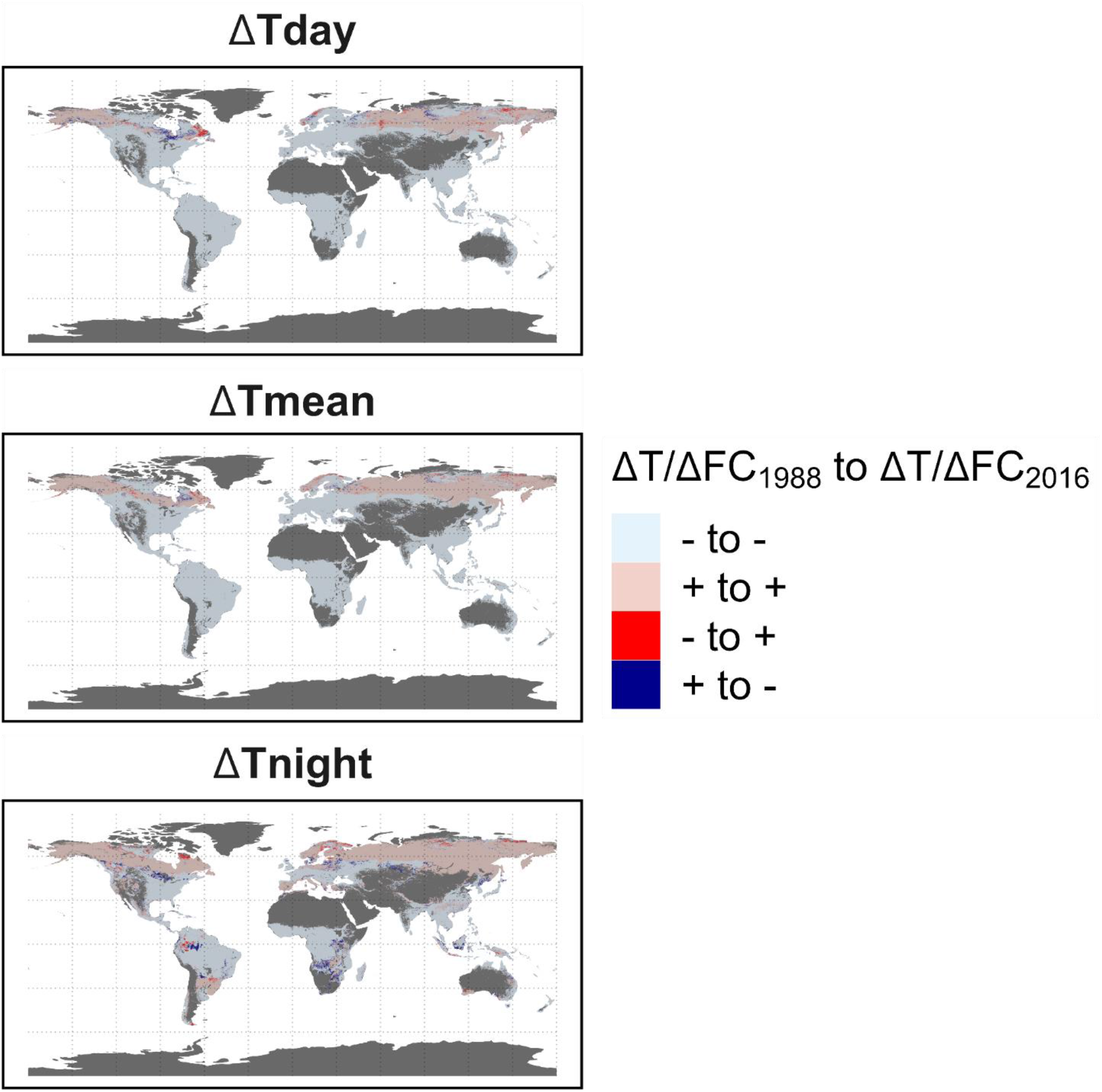
Change in the sign of the day (ΔTday), mean (ΔTmean), and night temperature (ΔTnight) sensitivity to forest cover change between the years 1988 and 2016.

**Fig. S16:**
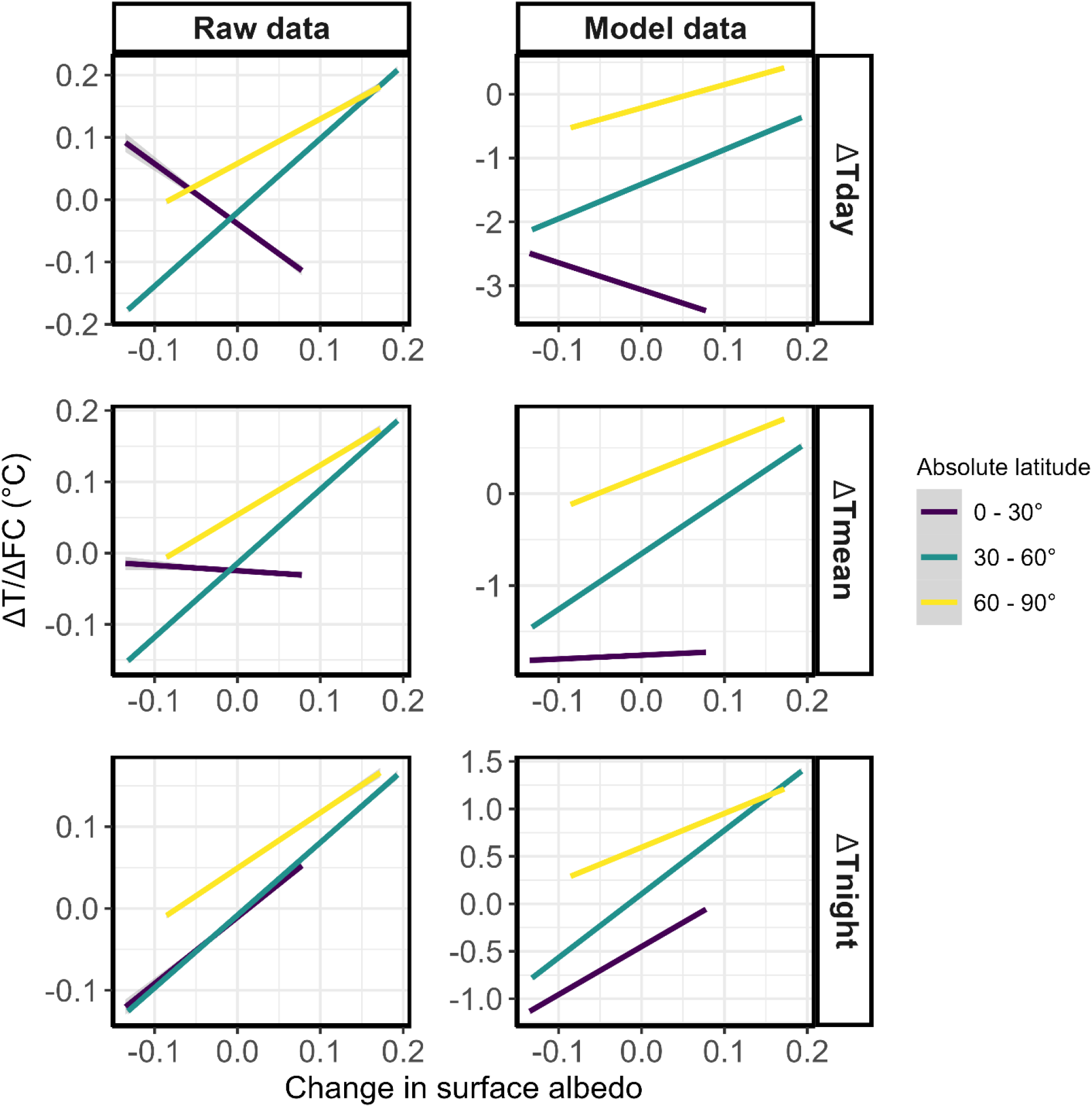
The relationship between the change in the surface albedo and sensitivity of land surface temperatures to forest cover change per 30°-absolute latitudinal interval. The raw data consist of observed changes in land surface temperatures divided by the corresponding forest cover change for the period 2003-2012 (n= 651,301). The model data represent the average predicted sensitivities of land surface temperatures to forest cover during the same period, at locations with significant forest cover change (n= 651,301). The change in surface albedo is considered for the same period. All linear regression lines were significant (P-values >0.05).

## Extended Tables

**Table S1:** Overview of predictor variables used for clustering.

| Variables | Sources |
| --- | --- |
| CHELSA_BIO_Annual_Mean_Temperature | http://chelsa-climate.org/bioclim/ |
| CHELSA_BIO_Annual_Precipitation | http://chelsa-climate.org/bioclim/ |
| CHELSA_BIO_Isothermality | http://chelsa-climate.org/bioclim/ |
| CHELSA_BIO_Max_Temperature_of_Warmest_Month | http://chelsa-climate.org/bioclim/ |
| CHELSA_BIO_Mean_Diurnal_Range | http://chelsa-climate.org/bioclim/ |
| CHELSA_BIO_Mean_Temperature_of_Coldest_Quarter | http://chelsa-climate.org/bioclim/ |
| CHELSA_BIO_Mean_Temperature_of_Driest_Quarter | http://chelsa-climate.org/bioclim/ |
| CHELSA_BIO_Mean_Temperature_of_Warmest_Quarter | http://chelsa-climate.org/bioclim/ |
| CHELSA_BIO_Mean_Temperature_of_Wettest_Quarter | http://chelsa-climate.org/bioclim/ |
| CHELSA_BIO_Min_Temperature_of_Coldest_Month | http://chelsa-climate.org/bioclim/ |
| CHELSA_BIO_Precipitation_of_Coldest_Quarter | http://chelsa-climate.org/bioclim/ |
| CHELSA_BIO_Precipitation_of_Driest_Month | http://chelsa-climate.org/bioclim/ |
| CHELSA_BIO_Precipitation_of_Driest_Quarter | http://chelsa-climate.org/bioclim/ |
| CHELSA_BIO_Precipitation_of_Warmest_Quarter | http://chelsa-climate.org/bioclim/ |
| CHELSA_BIO_Precipitation_of_Wettest_Month | http://chelsa-climate.org/bioclim/ |
| CHELSA_BIO_Precipitation_of_Wettest_Quarter | http://chelsa-climate.org/bioclim/ |
| CHELSA_BIO_Precipitation_Seasonality | http://chelsa-climate.org/bioclim/ |
| CHELSA_BIO_Temperature_Annual_Range | http://chelsa-climate.org/bioclim/ |
| CHELSA_BIO_Temperature_Seasonality | http://chelsa-climate.org/bioclim/ |
| ESA_SumSnowWaterEquivalent | https://opensciencedata.esa.int/variables/snow-water-equivalent |
| EarthEnvTopoMed_Slope | https://www.earthenv.org/topography |
| EarthEnvTopoMed_VectorRuggednessMeasure | https://www.earthenv.org/topography |
| EarthEnvTopoMed_Elevation | https://www.earthenv.org/topography |
| EarthEnvTopoMed_Roughness | https://www.earthenv.org/topography |
| SG_Clay_Content_015cm | https://www.soilgrids.org ;<br>https://www.isric.org/explore/wosis/accessingwosis-derived-datasets |
| SG_Coarse_fragments_015cm | https://www.soilgrids.org ;<br>https://www.isric.org/explore/wosis/accessingwosis-derived-datasets |
| SG_Depth_to_bedrock | https://www.soilgrids.org ;<br>https://www.isric.org/explore/wosis/accessingwosis-derived-datasets |
| SG_H2O_Capacity_015cm | https://www.soilgrids.org ;<br>https://www.isric.org/explore/wosis/accessingwosis-derived-datasets |
| SG_H2O_Capacity_030cm | https://www.soilgrids.org ;<br>https://www.isric.org/explore/wosis/accessingwosis-derived-datasets |
| SG_Sand_Content_015cm | https://www.soilgrids.org ;<br>https://www.isric.org/explore/wosis/accessingwosis-derived-datasets |
| SG_Silt_Content_015cm | https://www.soilgrids.org ;<br>https://www.isric.org/explore/wosis/accessingwosis-derived-datasets |

**Table S2:** The sensitivity of land surface temperatures and energy fluxes to forest cover change. The coefficient and standard error of the coefficient of forest cover change (°C per 100% forest restoration) in the biome-level multivariate linear models for the period 2003-2012. The R-squared of the models is shown between brackets.

| Dependent variable | Boreal<br>(n=195,892) | Temperate<br>(n=158,851) | Arid<br>(n=17,598) | Tropical<br>(n=278,587) |
| --- | --- | --- | --- | --- |
| $\Delta T_{day}$ | $-0.02 \pm 0.01$<br>(0.13) | $-2.15 \pm 0.02$<br>(0.27) | $-5.51 \pm 0.10$<br>(0.35) | $-2.96 \pm 0.02$<br>(0.22) |
| $\Delta T_{mean}$ | $0.42 \pm 0.01$<br>(0.16) | $-1.32 \pm 0.01$<br>(0.34) | $-2.81 \pm 0.05$<br>(0.31) | $-1.68 \pm 0.01$<br>(0.19) |
| $\Delta T_{night}$ | $0.85 \pm 0.01$<br>(0.19) | $-0.50 \pm 0.02$<br>(0.33) | $-0.10 \pm 0.03$<br>(0.03) | $-0.40 \pm 0.01$<br>(0.03) |
| $\Delta SW$ | $2.40 \pm 0.01$<br>(0.47) | $2.03 \pm 0.01$<br>(0.38) | $1.82 \pm 0.04$<br>(0.38) | $1.07 \pm 0.01$<br>(0.39) |
| $\Delta LE$ | $-0.95 \pm 0.01$<br>(0.21) | $-1.01 \pm 0.02$<br>(0.16) | $-4.34 \pm 0.11$<br>(0.24) | $-2.27 \pm 0.02$<br>(0.08) |
| $\Delta GH$ | $-0.90 \pm 0.01$<br>(0.12) | $-2.08 \pm 0.02$<br>(0.19) | $0.23 \pm 0.10$<br>(0.13) | $-0.28 \pm 0.03$<br>(0.05) |

## Notes

### Competing Interest Statement

The authors have declared no competing interest.

